# MLKS2 is an ARM domain and F-actin-associated KASH protein that functions in stomatal complex development and meiotic chromosome segregation

**DOI:** 10.1101/609594

**Authors:** Hardeep K. Gumber, Joseph F. McKenna, Andrea F. Tolmie, Alexis M. Jalovec, Andre C. Kartick, Katja Graumann, Hank W. Bass

## Abstract

The linker of nucleoskeleton and cytoskeleton (LINC) complex is an essential multi-protein structure spanning the eukaryotic nuclear envelope. The LINC complex functions to maintain nuclear architecture, positioning, and mobility, along with specialized functions in meiotic prophase and chromosome segregation. Members of the LINC complex were recently identified in maize, an important scientific and agricultural grass species. Here we characterized *Maize LINC KASH AtSINE-like2*, *MLKS2*, which encodes a highly conserved SINE-group plant KASH protein with characteristic N-terminal armadillo repeats (ARM). Using a heterologous expression system, we showed that actively expressed GFP-MLKS2 is targeted to the nuclear periphery and colocalizes with F-actin and the endoplasmic reticulum, but not microtubules in the cell cortex. Expression of GFP-MLKS2, but not GFP-MLKS2ΔARM, resulted in nuclear anchoring. Genetic analysis of transposon-insertion mutations, *mlks2-1* and *mlks2-2*, showed that the mutant phenotypes were pleiotropic, affecting root hair nuclear morphology, stomatal complex development, multiple aspects of meiosis, and pollen viability. In male meiosis, the mutants showed defects for bouquet-stage telomere clustering, nuclear repositioning, perinuclear actin accumulation, dispersal of late prophase bivalents, and meiotic chromosome segregation. These findings support a model in which the nucleus is connected to cytoskeletal F-actin through the ARM-domain, predicted alpha solenoid structure of MLKS2. Functional conservation of MLKS2 was demonstrated through genetic rescue of the misshapen nuclear phenotype of an Arabidopsis (triple-*WIP*) KASH mutant. This study establishes a role for the SINE-type KASH proteins in affecting the dynamic nuclear phenomena required for normal plant growth and fertility.

## Introduction

The cell nucleus is a eukaryotic organelle that houses, organizes and expresses the bi-parentally inherited genetic material. As a cellular compartment, the nucleus is separated from the cytoplasm by the multifunctional double-membraned nuclear envelope (NE). The NE has embedded macromolecular protein complexes, such as the nuclear pore complex (NPC) and the linker of nucleoskeleton and cytoskeleton (LINC) complex, which coordinate fundamental cell processes such as signaling and gene regulation (reviewed by Hetzer, 2010; Van de Vosse et al., 2011; D’Angelo, 2018).

The LINC complex was discovered relatively recently, but has been shown to be highly conserved in eukaryotes, with myriad functions as described in numerous reviews (Burke and Roux, 2009; Chang et al., 2015; Hieda, 2017; Kim et al., 2015; Luxton and Starr, 2014; Meier et al., 2017). These LINC-mediated functions include regulation of nuclear shape and position (Dittmer and Richards, 2008; Sakamoto and Takagi, 2013), mechanosensory signal transduction (Uzer et al., 2016), control of cell division, intranuclear architecture and gene regulation (Alam et al., 2016; Poulet et al., 2017b; Wang et al., 2018), and the specialized behavior of meiotic chromosomes to ensure segregation and fertility (Link et al., 2014; Murphy et al., 2014; Varas et al., 2015). The two core LINC components are the inner nuclear membrane (INM) SUN-domain proteins (Hagan and Yanagida, 1995; Malone et al., 1999) and outer nuclear membrane (ONM) KASH-domain proteins (Starr and Han, 2002). The SUN domain proteins are relatively well conserved across taxa and can be recognized by sequence homology and domain composition which includes transmembrane domains, the SUN domain, and a coiled-coil domain. In contrast, KASH proteins are functionally conserved, with a C-terminal region that includes a transmembrane domain and diagnostic terminal residues, but otherwise can be quite divergent with limited apparent homology (Zhou and Meier, 2013; Evans et al., 2014; Kim et al., 2015; Meier, 2016; Poulet et al., 2017a).

Structural studies reveal that SUN proteins form trimers that can bind to three KASH proteins (Sosa et al., 2012). The cytoplasmic N-terminal regions of KASH domain proteins interact directly or indirectly with a variety of cytoskeletal structures including motor proteins, microfilaments, microtubules, or intermediate filaments (Starr and Fridolfsson, 2010; Tamura et al., 2013; Luxton and Starr, 2014; Varas et al., 2015). The forces generated by the cytoskeletal elements result in LINC-dependent movements known or thought to be required for specialized nuclear morphology associated with mitotic and meiotic cell division, karyogamy at fertilization, cell polarity, and response to biotic and abiotic interactions (Starr and Fridolfsson, 2010; Gundersen and Worman, 2013; Griffis et al., 2014; Fernández-Álvarez et al., 2016; Groves et al., 2018).

Most of our knowledge of the LINC complex comes from studies of metazoans and yeast. However, given the global challenges in agriculture and the need for plant-based biorenewable resources (Godfray et al., 2010), it is important to better understand the plant NE and its fundamental components. Recently, progress towards understanding plant LINC biology has expanded beyond Arabidopsis to include Medicago and maize (Gumber et al., 2019; Newman-Griffis et al., 2019).

The first SUN domain protein identified in plants was OsSad1, found in a nuclear proteomic study in rice (Moriguchi et al., 2005). Subsequently, SUN domain proteins have been identified in many plant species (reviewed by Meier, 2016; Meier et al., 2017; Poulet et al., 2017a) with roles in the maintenance of nuclear shape and size (Graumann et al., 2010; Oda and Fukuda, 2011; Graumann et al., 2014) and meiotic chromosome behavior (Murphy et al., 2014; Varas et al., 2015). In maize, SUN2 has been shown to form a novel structure called the meiotic SUN belt during prophase of meiosis I (Murphy et al., 2014). The SUN belt region of the NE includes the telomere cluster which defines the zygotene bouquet (Murphy et al., 2014).

The first plant KASH proteins were originally discovered in Arabidopsis through analysis of RanGAP-NE anchoring proteins, and implicated in NE-associated signaling and maintenance of nuclear shape (Xu et al., 2007; Zhou et al., 2012). They were later shown to bind to SUN (Meier et al., 2010; Zhou et al., 2012) and possess a single near-C-terminal TMD and a C-terminal amino acid sequence of V/P-P-T. These features are diagnostic of plant KASH-like proteins and were collectively used as criteria for identification of ten KASH candidates in maize (Gumber et al., 2019). From that study, four groups of <u>M</u>aize <u>L</u>INC <u>K</u>ASH (MLK) related genes were identified: the *AtWI<u>P</u>*-like group (*MLKP1* - *MLKP4*), the *At<u>S</u>INE*-like group (*MLKS1* and *MLKS2*), the <u>g</u>rass-specific group (*MLKG1* and *MLKG2*), and the KASH-associated *AtWI<u>T</u>*-like group (*MLKT1* and *MLKT2*).

A particularly interesting and distinct group of plant KASH-like proteins are the SUN-interacting nuclear envelope (SINE) proteins, which contain armadillo domains (Armadillo-type fold, InterPro ID: IPR000225). The armadillo (ARM) type fold is a protein-protein interaction surface formed by twisting of multiple alpha helices around a central axis, and has been reported in a variety of proteins involved in intracellular signalling and cytoskeletal regulation (Coates, 2003; Hatzfeld, 1998). Genetic analysis of the SINE-group KASH genes in Arabidopsis reveals non-redundant functions ranging from actin-dependent nuclear positioning in guard cells for *AtSINE1* to innate immunity against a fungal pathogen for *AtSINE2* (Zhou et al., 2014). Despite their deep conservation, the biological functions of plant SINE-group KASH genes in monocot grass species remain largely uncharacterized. Maize contains two *AtSINE1* homologs, *MLKS1* (Zm00001d031134) and *MLKS2* (Zm00001d052955). Both of these contain an ARM domain towards their N-terminus (Gumber et al., 2019). Given the fundamental importance of connecting the nucleus to the cytoplasm, these maize SINE-group KASH genes were predicted to participate in a broad array of processes requiring coordinated actions of the NE with the cytoskeleton.

In this study, we have undertaken an investigation of a single maize LINC gene, *MLKS2*, for several reasons. First of all, the MLKS2 protein was detected in CoIP assays using ZmSUN2 antisera with earshoot nuclei (Gumber et al., 2019). Secondly, *MLKS2* is expressed in actively dividing cells, including meiosis-stage tassels, according to transcriptomic studies (Stelpflug et al., 2016). Thirdly, *MLKS2* orthologs are conserved across land plants, increasing the potential translational impact of the study (Gumber et al., 2019). Finally, two independent Mu transposon insertion alleles of *MLKS2* were available (https://maizegdb.org/uniformmu) allowing for genetic and phenotypic analyses. Here we characterize *MLKS2* using genetic and cellular experiments in order to: (1) test for canonical KASH properties of NE localization and SUN-binding; (2) investigate colocalization with cytoplasmic structures; (3) address its role in meiosis and nuclear shape and positioning via mutant phenotyping; (4) gain mechanistic insight into its role in nuclear positioning and actin interaction using domain deletion assays; and (5) test for functional conservation through cross-species genetic rescue of nuclear shape phenotype.

## Results

The structure of the *MLKS2* gene and protein are summarized in Figure 1. The *MLKS2* gene Zm00001d052955_T001 (from B73 v4) has three exons and is predicted to encode a protein with 637 amino acids (Fig. 1A-B). The secondary and domain structure of the MLKS2 protein includes the canonical KASH features of a single transmembrane domain near the C-terminus, followed by a 14-residue C-terminal KASH domain and terminal sequence of LVPT (Fig. 1C). Nearly half of the protein (8-294) has an armadillo-type fold domain (InterPro IPR016024). Another large segment (301-598) is composed of three segments classified as disordered regions, designated here as DR1 (301-401), DR2 (414-478), and DR3 (488-598). Tertiary structure prediction using I-TASSER revealed that the ARM and disordered regions could adopt a single large structure similar to that of human protein phosphatase 2A, PP2A, a structural scaffolding subunit. PP2A (pdb 1b3uA) was the top ranked structural analog as shown separately and threaded together with MLKS2 (Fig. 1D). This overall structure is referred to as an alpha solenoid, broadly deployed in biological systems as a binding structure (Fournier et al., 2013).

**Figure 1:**
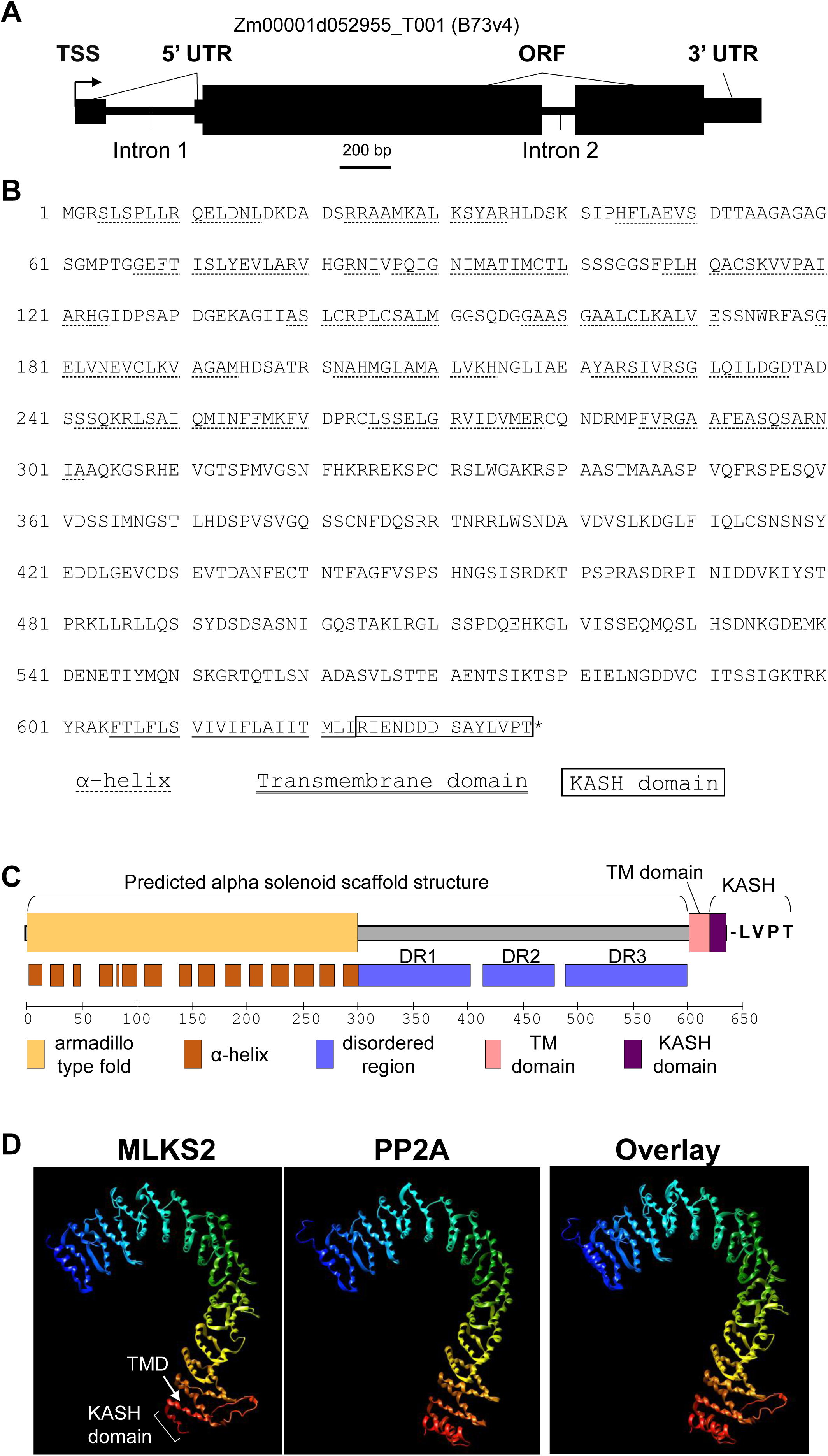
Gene model and protein domain diagram of *MLKS2*. A) Gene model diagram for *MLKS2* with gene features drawn to scale. Total length of the primary transcript (transcript model for B73v4 is indicated above) is 3007 bp. Three exons are shown as black boxes. The 5’UTR of the gene is interrupted by “Intron 1.” Gene features are indicated: TSS, transcription start site; UTR, untranslated region; ORF, open reading frame. B) The amino acid sequence of MLKS2 with domains underlined with lines of different styles, or boxed. The protein sequence is identical to that from genotype W22 (not shown) used for transposon mutagenesis. The key is given under the sequence. C) Domain diagram for MLKS2 showing the names and locations of secondary structural features (alpha helices) or domains are indicated. The conserved terminal four amino residues (LVPT) at the end of the KASH domain are shown. D) The I-TASSER protein structure prediction software was used with MLKS2 as input sequence. The top-ranked structural homolog is shown for a predicted tertiary structure of MLKS2 (1st panel) next to the known structure of PP2A structural subunit (pdb 1b3uA, 2nd panel), followed by an overlay of the two (3rd panel). The rainbow annotation denotes the protein polarity from the N-terminus (blue) to the C-terminus (red). The transmembrane (TMD) and KASH domains of MLKS2 are annotated.

### Nuclear envelope localization of MLKS2 is disrupted by C-terminal but not ARM domain deletions

We first examined the cellular localization of MLKS2 and its potential to interact with SUN or the cytoskeleton. GFP-tagged MLKS2 was transiently expressed and imaged as shown in Figure 2, using the tobacco expression system previously described (Gumber et al., 2019). The GFP-MLKS2 localized to the nuclear periphery as expected for a KASH protein (Fig. 2B). The GFP-MLKS2 was also seen at the cell periphery in a conspicuous pattern resembling that of a cytoskeletal network (Fig. 2B). In order to map the protein regions responsible for these patterns, we made a series of domain deletion constructs (Fig. 2A). The deletions were MLKS2Δ4, with removal of the terminal four amino acid residues; MLKS2ΔKASH, with replacement of the terminal KASH domain with three alanine residues; MLKS2ΔTM, with removal of the transmembrane domain; or MLKS2ΔARM, with removal of the entire Armadillo fold region, DR1, DR2, and part of DR3.

**Figure 2:**
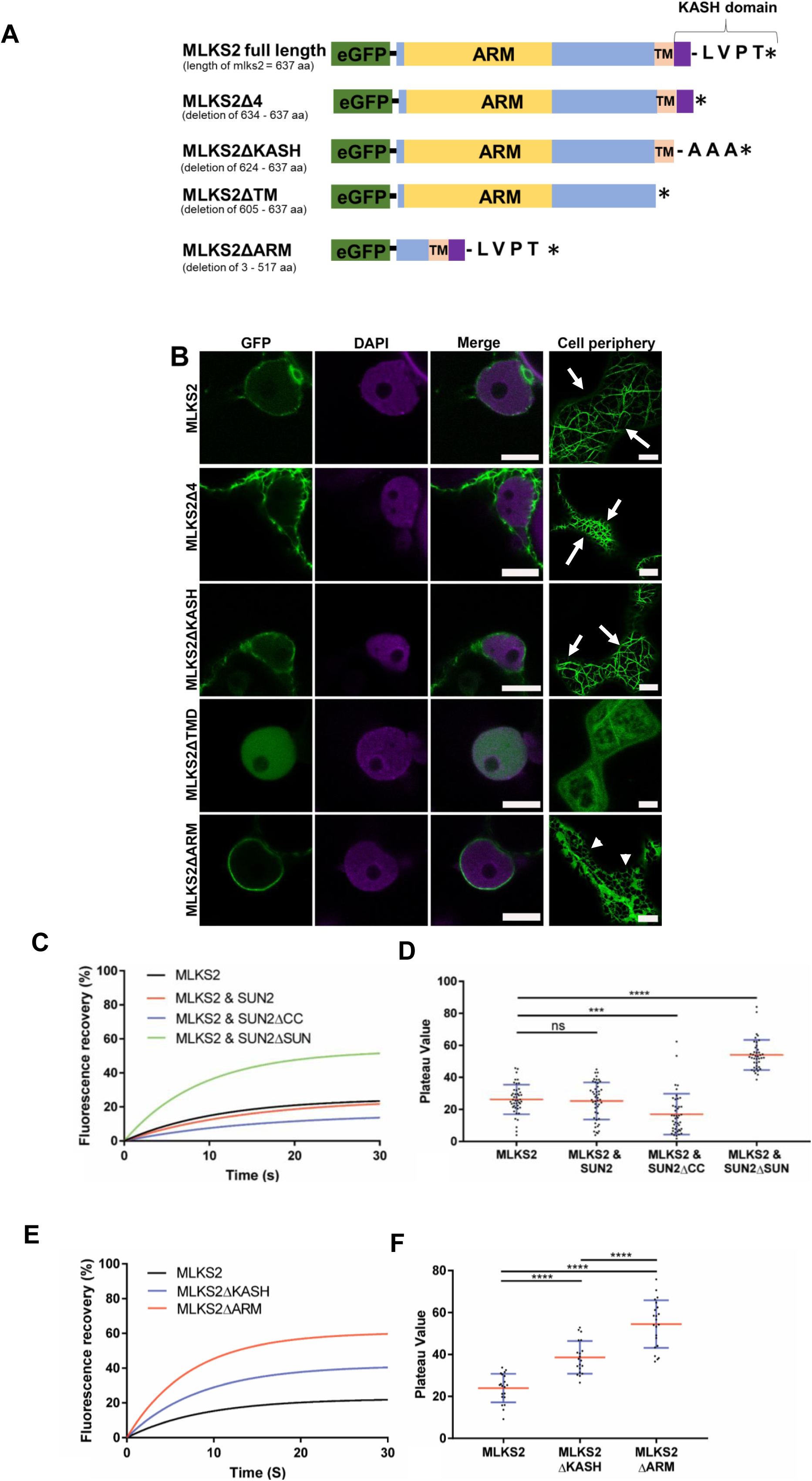
MLKS2 cellular localization and ZmSUN2-interaction. Subcellular localization of full length and deletion mutants of GFP-MLKS2 in transiently expressing *N. benthamiana* leaf cells as previously described (Gumber et al., 2019). A) Domain deletion diagram of all constructs used showing the ARM domain (yellow), disordered regions (blue), transmembrane domain (TM), KASH domain (purple), terminal AA residues, and stop codon (*). B) Full length GFP-MLKS2 (green) and all but one deletion mutant localize to the NE around DAPI-stained chromatin (magenta). GFP-MLKS2ΔTM appears soluble and distributed throughout the nucleoplasm. A cell periphery network-like pattern was observed for all constructs, with GFP-MLKS2ΔTM appearing as soluble in the cytoplasm and GFP-MLKS2ΔARM appearing as associated with the ER. C) Fluorescence recovery curve from MLKS2 FRAP alone or when co-expressed with mCherry-ZmSUN2, mCherry-ZmSUN2ΔCC (without coiled coil domain), or mCherry-ZmSUN2ΔSUN (without SUN domain). D) FRAP recovery plateau values for fluorescence recovery curves shown in C). E) Fluorescence recovery curves showing increased mobile fraction in the NE for KASH or ARM domain deletion variants of GFP-MLKS2. F) FRAP recovery plateau values of MLKS2, MLKS2ΔKASH, and MLKS2ΔARM shown in D). Scale bar denotes 10 µm. For whisker plots, blue lines denote SD error bars, red lines denote mean. Nuclei (N = ≥30) imaged across three experimental repeats per treatment. ANOVA statistical test used where Ns = P≥0.05, *** = P≤0.001 and **** = P≤0.0001.

We found that the terminal Δ4 and ΔKASH constructs reduced the relative abundance of MLKS2 at the nuclear periphery but retained the network-like pattern in cytoplasm as clearly seen at the cell periphery (arrows, Fig. 2B). In contrast, removal of the TM domain completely changed the localization pattern at both the nuclear and the cell periphery to one resembling that of a soluble protein showing nucleoplasmic and cytoplasmic labelling. The N-terminal deletion construct, ΔARM, retained its NE localization but with a more uniform and less patchy signal. However, the ΔARM variant had a dramatic effect on the cell peripheral labelling, which changed from a filamentous cytoskeleton-like pattern to a more endoplasmic reticulum-like pattern (arrow heads, Fig. 2B), including both sheets and tubules. These results demonstrate that when expressed under the control of a constitutive promoter, MLKS2 can accumulate at two places in the cell, the NE requiring C-terminal domains and the cytoplasm requiring the N-terminal ARM domains. It remains a formal possibility that the non-nuclear cytoplasmic MLKS2 may be an artifact of experimental over- or heterologous expression and may not, therefore, reflect its primary biological function.

### The KASH and ARM domains immobilize the NE-associated MLKS2

We next tested for interaction of MLKS2 with ZmSUN2 using fluorescence recovery after photobleaching (FRAP). The mobility of GFP-MLKS2 expressed alone was compared to that co-expressed with full length mCherry-ZmSUN2, or deletion constructs lacking the SUN domain or coiled-coil regions of ZmSUN2 (Fig. 2C, D, S1). GFP-MLKS2 has a large immobile fraction to start with, showing only 25% fluorescence recovery 30 s after photobleaching (Fig. 2D). When co-expressed with full length SUN2, the immobile fraction of GFP-MLKS2 showed a slight but statistically insignificant increase compared to GFP-MLKS2 alone (Fig. 2C, D). When co-expressed with SUN2 protein lacking the coiled-coil domain (SUN2ΔCC), the immobile fraction of GFP-MLKS2 was significantly increased compared to GFP-MLKS2 alone or when co-expressed with full length ZmSUN2. When co-expressed with ZmSUN2 protein lacking the SUN domain (SUN2ΔSUN), the immobile fraction of GFP-MLKS2 was significantly and markedly decreased compared to GFP-MLKS2 alone or when co-expressed with full length ZmSUN2.

Given the large immobile fraction of MLKS2 to begin with, we did not expect to observe a marked increase in its immobile fraction when co-expressed with full length SUN. However, the drastic reduction in the immobile fraction when co-expressed with SUN2ΔSUN, compared to co-expression with full-length SUN, was unexpected but consistent with the interpretation that MLKS2 interacts with SUN2 via its SUN domain. One possible explanation for the unexpected effect of the SUN2ΔSUN is that its expression could have a dominant negative effect on the tobacco nuclear envelope components and their ability to interact with GFP-MLKS2. We conclude that two regions of MLKS2 each play a role in anchoring it within the NE, the C-terminal KASH domain and the N-terminal ARM domain.

We used the FRAP mobility assay to decipher the contributions of the KASH and ARM domains to the relatively large immobile fraction of MLKS2. Removal of the KASH domain decreased the immobile fraction (Fig. 2E, F, S1), consistent with a KASH-SUN binding interaction in the NE. Remarkably, removal of the ARM domain decreased the immobile fraction even more (Fig. 2E, F) making over 60% of the protein population mobile. This indicates that SINE2-type KASH proteins are likely docked in the NE through interactions at both ends, one via SUN-KASH binding in the NE and one via ARM domain binding to the cytoskeleton.

### Genetic analysis of two transposon-disrupted alleles of *MLKS2*

For genetic analysis, we searched the UniformMu transposon mutagenesis records (McCarty and Meeley, 2009) at maizeGDB and found two alleles for *MLKS2*. The mutator element mobilized in UniformMu is MuDR, a 1.4 kbp non-autonomous *Mu1* element (Barker et al., 1984). The two insertion alleles are designated here as *mlks2-1* and *mlks2-2*, as diagrammed in Figure 3. Both alleles involved insertions into the protein coding region in Exon 2. PCR primers were selected to confirm the location and sequences surrounding the transposon insertion sites (Fig. 3 A,B). For each allele, a 9-bp target site duplication was detected (underlined, Fig. 3B), characteristic for *Mu* insertions. The PCR reactions used for genotyping of individual plants are shown for representative individuals from families segregating for the insertions (Fig. 3C)

**Figure 3.**
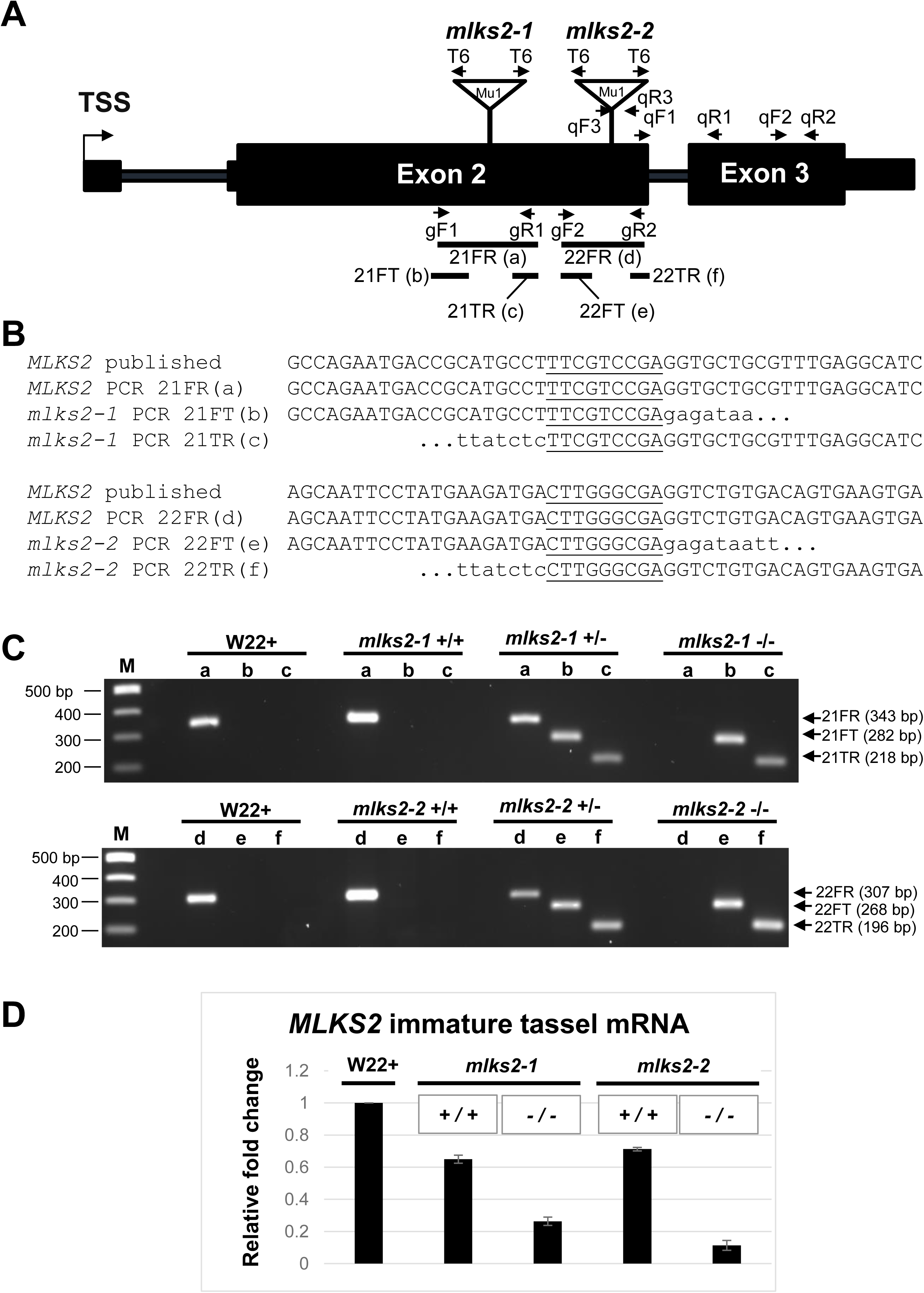
Genotypic characterization of *MLKS2* alleles. Two transposon-tagged alleles of *MLKS2, mlks2-1* and *mlks2-2* are depicted. A) *MLKS2* gene model showing the location of *Mu1* transposon insertion sites (triangles) for two alleles. The positions of allele-specific primer pairs (gF1, gR1 and gF2, gR2) and Tir6 primer (T6) at the Mu1 transposon terminal inverted repeat sequence are marked with arrows. The positions of primer pairs (qF1, qR1; qF2, qR2; or qF3, qR3) used for qRT-PCR are also marked with arrows. The PCR products (21FR, 21FT, 21TR, 22FR, 22FT, and 22TF) used for sequence verification, genotyping, and quantitative RT-PCR are indicated below the gene model. B) Sequences aligned around the insertion site include the published parental sequence (Springer et al., 2018), the wild-type allele from PCR products (“a” and “d”) from W22, and the mutator-flanking sequences from PCR products (“b”,“c”, “e”, and “f”) using one gene primer and one mutator-specific primer (T6). In both alleles, a 9-bp duplication (underlined) was detected. C) Agarose gels showing PCR products amplified from W22 and plants from families segregating for *mlks2-1* (top gel) or *mlks2-2* (bottom gel) allele. The PCR products were amplified using gene-specific primer pairs (lanes/PCR products a, d) or primer pairs from one gene-specific primer and one mutator (T6) primer (lanes/PCR products b, c, e, f). Plant genotypes are shown on the top of the gels, primer pairs and band sizes are indicated on the right. The lanes “M” contain 100 bp DNA marker fragments at the lengths indicated. D) Fold change in the transcript levels of *MLKS2* in families segregating for *mlks2-1* or *mlks2-2*; homozygous WT siblings (+/+) or homozygous mutant plants (−/−) were quantified relative to W22 using an average of 3 primer pairs (qF1-qR1, qF2-qR2, qF3-qR3) as measured by qRT-PCR.

We examined RNA from immature, meiosis-staged tassels of wild-type and mutant plants to check if the transposon insertions reduced the mRNA levels. Quantitative RT-PCR showed that both alleles, *mlks2-1* and *mlks2-2*, resulted in decreased relative RNA levels when compared to those from wild-type siblings (*Mu −/−*) of *mlks2-1* and *mlks2-2* or those from parental W22 line of maize (Fig. 3D). Of the two alleles, *mlks2-2* showed the greatest reduction in transcript levels.

### *MLKS2* is required for normal male meiosis and pollen development

Given the role of the LINC complex and NE in meiosis, we asked whether genetic disruption of *MLKS2* resulted in meiotic or post-meiotic phenotypes. In maize, like most eukaryotes, the NE and the LINC complex have highly specialized functions including a role in telomere bouquet formation, pairing, synapsis, and recombination, all of which must occur with high fidelity to ensure proper genome reduction and euploid transmission. We therefore used 3D cytology to look for several hallmarks of normal male meiosis, including the telomere bouquet, the eccentric positioning of the nucleus at early prophase I, 10 bivalents at late prophase I indicative of complete pairing, chiasma formation, and equal distribution of chromosomes after meiosis I and II as evidenced by daughter nuclei with matching sizes and shapes.

The *mlks2-2* mutants showed deviations from all of the cytological meiotic hallmarks inspected as summarized in Figure 4. At early meiotic prophase, 3D telomere FISH revealed that the *mlks2-2* plants showed partial telomere clustering on the NE at the bouquet stage (Fig. 4A-C) with variable numbers (4-12) of telomere FISH signals far removed from the primary cluster (bracketed “bq” in Fig. 4A, B). The magnitude of telomere dispersion was measured using pairwise telomere FISH signal distances per nucleus. The average pairwise telomere distance increased by 50% in the mutant, from 4.3 microns in normal nuclei (n=6) to 6.5 microns in *mlks2-2* nuclei (n=11). The tendency for multiple telomeres to reside far from the bouquet is also seen in the fold change of inter-telomere pairwise distances for the long distance bins (Fig. 4C). At early prophase I, meiotic nuclei tend to relocate towards one edge of the cell (Cowan et al., 2002), but *mlks2-2* nuclei remained closer to the cell center (Fig. 4D,E), according to measures of nuclear eccentricity. The cell-to-nucleus centroid offset at early prophase in normal cells decreased from 15 microns in normal (n=6) plants to 7 microns in mutant (n=11) plants (Fig. 4F).

**Figure 4:**
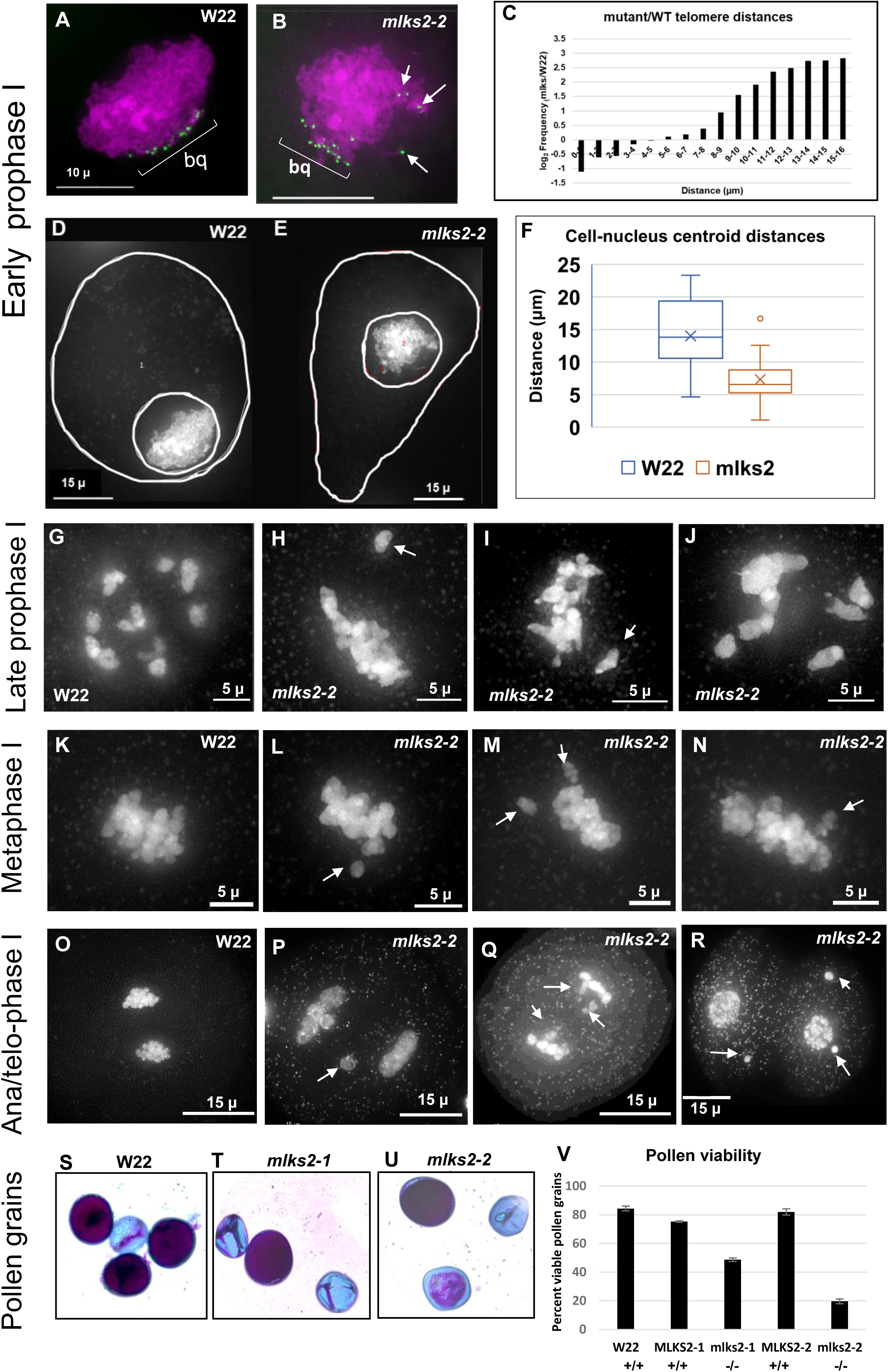
Multiple meiotic and post-meiotic defects of *mlks2* mutants. Cytological defects of *mlks2-2* during and after male meiosis. A) Early prophase I stage W22 meiocyte nucleus showing a typical bouquet (green dots, “bq”) of NE-associated telomeres visualized in the nucleus (DAPI, shown in magenta) using the 3D acrylamide oligo FISH method (Bass et al., 1997). B) Partial telomere bouquet in *mlks2-2* mutant at meiotic prophase, showing unusually distant telomeres (arrows) relative to the main bouquet (bq) telomere cluster region. C) Histogram showing bouquet-stage telomere pairwise distance distributions plotted as log2 fold change of mutant/wild-type per 1 micron distance bins (n=6 W22, n=11 *mlks2-2*). The mutants show a pronounced increase in the longer telomere-to-telomere distance bins. Nuclear position phenotypes for normal D) or mutant E) cells shown as projections from the middle-most 1/5 of the optical sections through the nuclei stained with DAPI, including traces around the cell and nuclear peripheries to ascertain 2D centroid locations. F) Eccentricity plots showing the distribution of distances of between the pairwise centroids of nuclei and cells for normal (W22) and mutant (*mlks2*) meiocytes at at the bouquet stage, using the same nuclei as those analyzed in A-C. G) Late prophase I stage W22 meiocyte showing bivalents spread throughout the volume of the nucleus. H-J) Late prophase I stage *mlks2-2* meiocytes showing clumping of bivalents. K) W22 meiocyte showing bivalents on a normal meiosis I metaphase plate. L-N) Mutant *mlks2-2* meiocytes showing one or more chromosomes (arrows) not located in the meiosis I metaphase plate. O) W22 meiocyte at late anaphase I or early telophase I. P-Q) Mutant *mlks2-2* late anaphase I or early telophase I showing irregularly positioned “laggard” chromosomes (arrows). R) Mutant mlks2-2 at telophase after meiosis I, before meiosis II, and showing micronuclei (arrows) that are associated with failure of chromosomes or chromosomal fragments to reach the spindle poles. S-U) Pollen viability stains for wild-type S) or mutant T, U) pollen. Dark purple indicates viable pollen, light blue indicates inviable pollen. V) Quantification of pollen viability with n=1,000 or more for each genotype.

In the late prophase I stages of diplotene or diakinesis, chromosomes continue to condense while successful disomic pairing and recombination becomes cytologically evident in the form of well-separated bivalents. These bivalents total up to the haploid chromosome number, 10 in the case of normal maize (Fig. 4G). Failure to properly pair or recombine typically results univalents or multivalents, which often fail to segregate properly at anaphase during meiosis I division. The *mlks2-2* mutation caused conspicuous and obvious departure from these normal late prophase I patterns (Fig. 4H-J). The meiotic chromosomes from mutant plants showed aggregation and pronounced clumping of multiple chromosomes at late prophase I, with relatively few clear examples of well separated bivalents (compare Fig. 4G to 4H-I). The tendency and degree of aggregation varied in the mutant plants, even for cells from the same anther. The majority of the mutant cells showed these late prophase I phenotypes. This late prophase I clumping phenotype is intriguing given that telomere-NE tethering normally persists even after the bouquet stage. These results place MLKS2 in the poorly understood pathway that ensures uniform dispersal of bivalents after pachytene.

Anaphase and telophase can be particularly informative stages for detection of abnormal chromosome segregation. We found that the *mlks2-2* mutants showed numerous such segregation defects, including chromosomes not in the metaphase I plate bundle (arrows, Fig 4L-N). The mutant plants frequently showed chromosome laggards (Fig. 4P-Q) and micronuclei (Fig. 4R), both of which are indicative of missegregation. Frequency of cells with one or more aberrant patterns at meiosis I (metaphase, anaphase, telophase) were low for wild-type (5%, n = 200) but high for *mlks2-2* (46%, n = 303) Specifically, the percentages of cells with abnormal chromosomes per stage were 61% in diakinesis (n=109), 37% in metaphase I (n=123), 60% in anaphase I (n=10), and 34% in the combined stages of telophase I, telophase II, and 4-nucleate tetrad (n=61).

We next checked for post-meiotic phenotypes associated with pollen viability. Acid fuchsin and malachite green differential pollen staining revealed that both *mlks2-1* and *mlks2-2* plants showed reduced pollen viability compared to normal plants (Fig. 4S-V, n>1,000 for each genotype). The pollen sterility phenotype was only partially penetrant and, like the other phenotypes, more severe for *mlks2-2*. Taken together, these phenotypes show that MLKS2 is required for proper male reproduction, with cytological phenotypes appearing as early as the zygotene bouquet stage of meiosis.

### *MLKS2* plays a conserved role in nuclear shape maintenance in root hair cells

Root hair nuclei in plants are pleomorphic, and in Arabidopsis their shape and positioning are disrupted by mutations in LINC or LINC-associated genes (Dittmer and Richards, 2008; Oda and Fukuda, 2011; Tamura et al., 2013; Zhou et al., 2012). Interestingly, this rounding up or loss of elongation nuclear shape phenotype appears to be generally diagnostic for genetic disruption of the LINC complex. We measured nucleus lengths in maize root hairs 5 days after imbibition, from normal and *mlks2* mutants. Normal (W22, also designated W22+ or wild-type) nuclei averaged a length of 37 microns, but mutant nuclei were only half that long on average (Fig. 5A, B). A related measure, circularity index from edge-traced nuclei in 2D images, also changed from 0.4 in normal plants to a more circular value of 0.7 in the more “rounded up” mutants (Fig. 5C). Although the biological role of root hair nuclear elongation remains to be determined, these shape measures provide a quick and quantitative phenotypic readout for genetic disruption of the LINC complex at the seedling stage.

**Figure 5.**
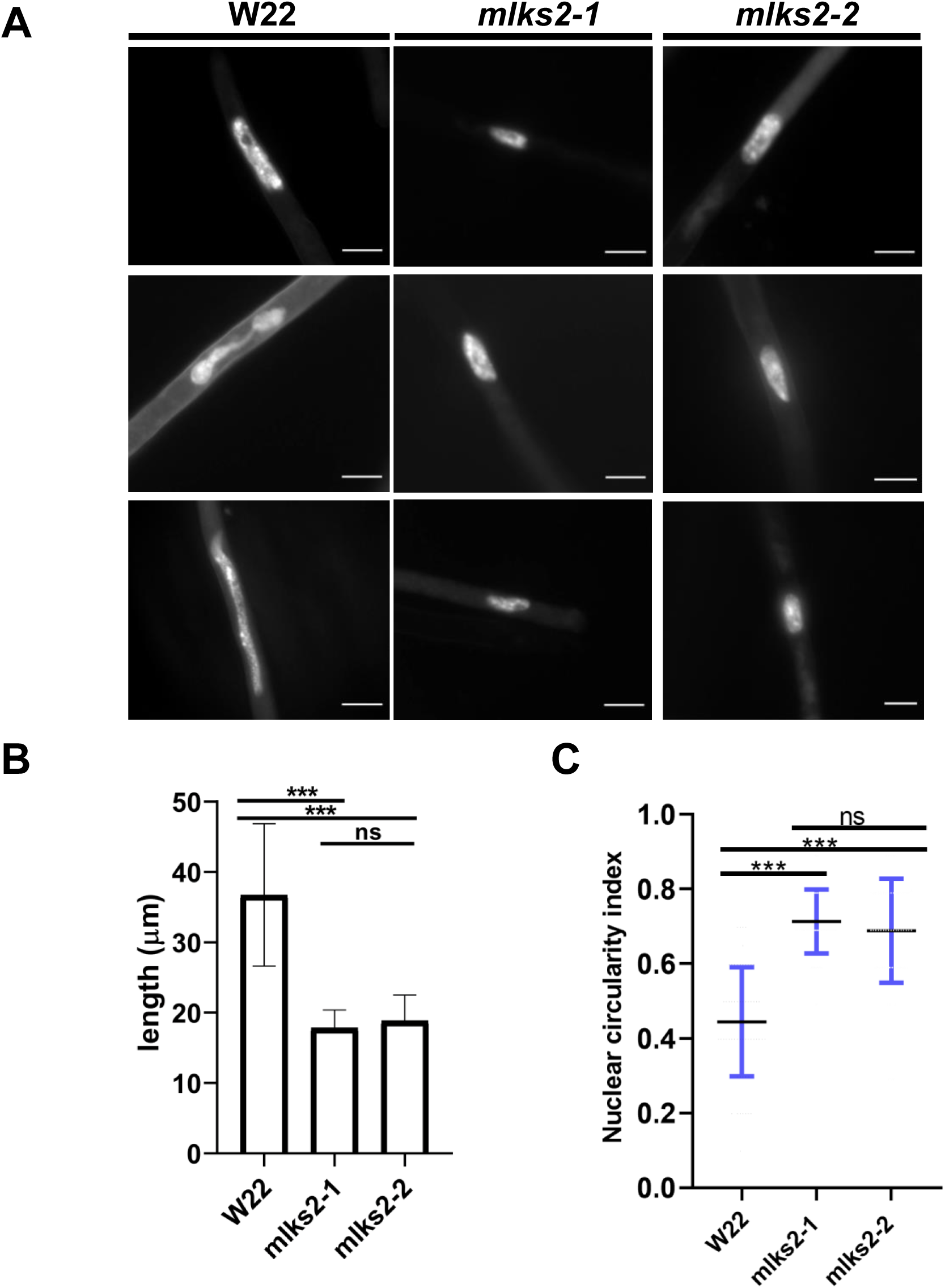
MLKS2 is required for the elongated nuclear shape phenotype in root hair cells. Root hair nuclei stained with DAPI in W22, *mlks2-1* and *mlks2-2* maize plants. B) Quantitation of longest dimension of W22, *mlks2-1*, or *mlks2-2* root hair nuclei. C) Circularity index measurements of nuclei from W22, *mlks2-1* and *mlks2-2* in which 1.0 represents a perfectly round shape.

### *MLKS2* is required for normal stomatal complex development

While screening seedling leaf tissues for nuclear shape phenotypes, we found an unexpected and dramatic effect on stomatal complex development in*mlks2* mutants, as summarized in Figure 6. Normal stomatal development in maize leaves (reviewed by Smith, 2001) results in a bilaterally symmetric 4-cell complex composed of 2 dumbbell-shaped guard cells (GC in Fig. 6A) flanked by two subsidiary cells (SC in Fig. 6A). In the *mlks2* mutants, the subsidiary cells, but not the guard cells, exhibited dramatic and variable deviations (denoted by arrows, Fig. 6D-I) in size, shape, and number per stomatal complex. Given the known role of asymmetric cell division in the normal development of the guard cell complex, we checked for cell polarity phenotypes at earlier stages of development. Images from the meristematic base of the fourth leaf showed that the *mlks2-2* mutant exhibited irregular cell polarities and division planes (Fig. 6L), abnormal shaped subsidiary cells (Fig. 6O), or extra inter-stomatal and subsidiary mother cells (Fig. 6K, N). These phenotypes closely resemble those of the *pangloss* or *brick* mutants of maize, known to perturb nuclear positioning and the actin cytoskeleton, culminating in irregular subsidiary cell development. These findings uncover an unanticipated developmental role for *MLKS2* that likely reflects its function in cell polarization and nuclear positioning for specialized, highly asymmetric cell divisions leading to the differentiation of the stomatal complex.

**Figure 6.**
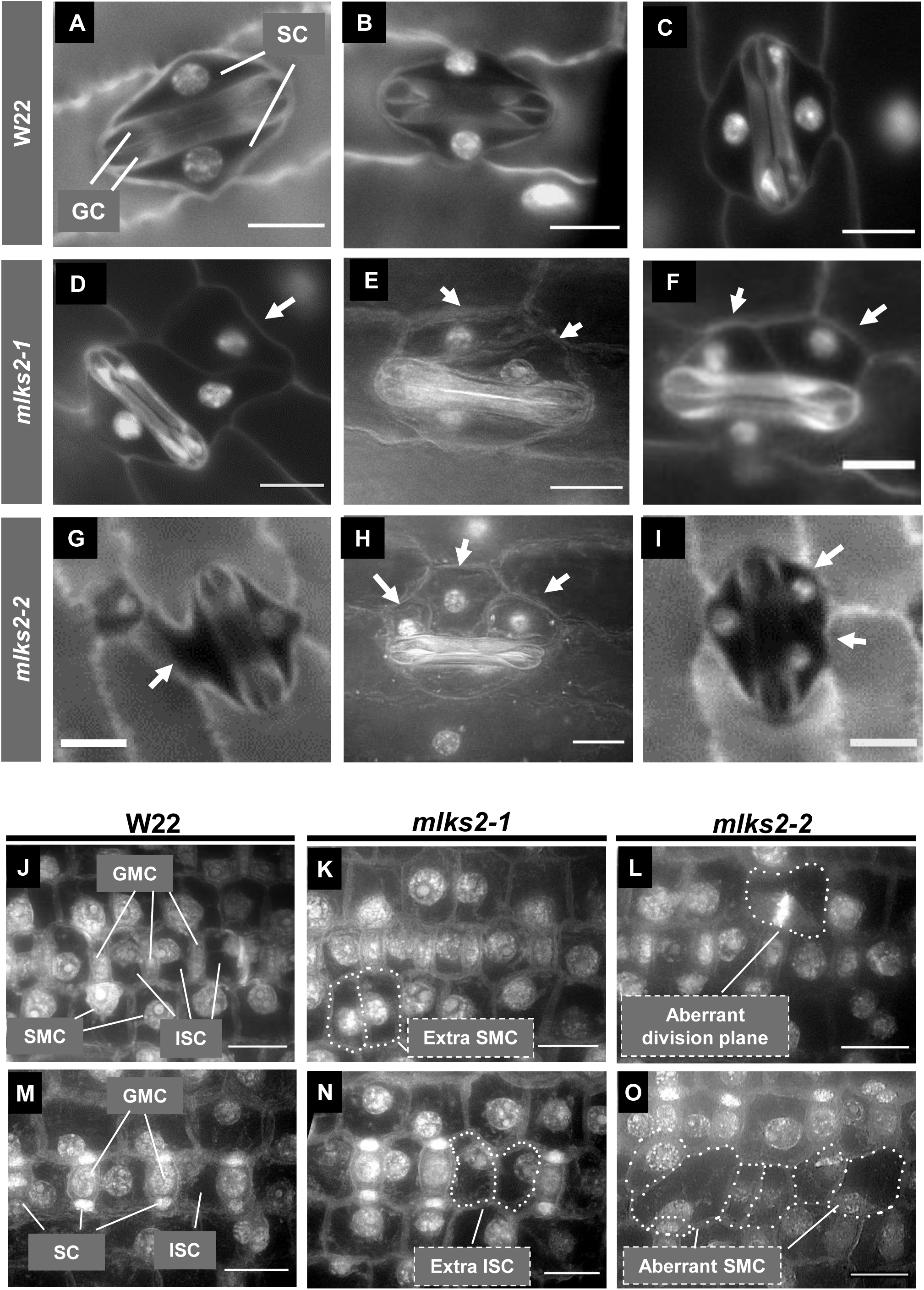
MLKS2 is required for normal stomatal complex development. Mature stomatal complex in DAPI stained leaf from wild-type W22 (A-C), *mlks2-1* (D-F) or *mlks2-2* (G-I) plants. For comparison, a typical stomatal complex has bilateral symmetry with two elongated central dumbbell-shaped guard cells (GC) flanked by two outer subsidiary cells (SC). In the *mlks2-1* D-F) or *mlks2-2* G-I) mutants, subsidiary cells appear abnormal (arrows) in their number, shape, or nuclear position relative to the guard cells. J-L) Representative images from early stages of stomatal development where subsidiary mother cells (SMC) are polarizing, with nuclei migrating towards the guard mother cells (GMC). Interstomatal cells (ISC) are annotated for W22. M-O) Representative images of developing stomatal complex after SMC polarization. Boundaries of abnormally shaped cells or cells with abnormal nuclear positioning are marked with dotted lines. Scale bars denote 15 µm.

### MLKS2 interacts with F-actin via its ARM domain and is required for recruitment of meiotic perinuclear actin

Given that MLKS2 has ARM domains, localizes to a cytoskeletal-like network, and shows mutant phenotypes suggestive of cytoskeletal interactions, we wanted to more directly explore the interaction of MLKS2 with actin. First, GFP-MLKS2 was co-expressed with markers for the endoplasmic reticulum (RFP-HDEL), the actin cytoskeleton (RFP-Lifeact) or the microtubule cytoskeleton (mCherry-TUA5) as shown in Figures 7, S2. Colocalization of GFP-MLKS2 was seen for both ER and F-actin but not microtubule markers (Fig. 7A, S2). The actin, but not the ER colocalization with MLKS2, requires the presence of the MLKS2 ARM domain region (Fig. 7B). To further characterize the interaction of MLKS2 with F-actin, we examined the localization of MLKS2 in cells treated with Latrunculin-B (LatB) to depolymerize the F-actin. This treatment caused the GFP-MLKS2 staining pattern to change from fibrous to a more ER-like network, similar to that of the ARM domain deletion of MLKS2 (compare 2B to 7C). These findings show that when expressed in tobacco, MLKS2 appears to induce an actin-ER colocalization. MLKS2 appears to connect the ER to F-actin through its ARM domain. Consistent with this, the mobile fraction of MLKS2 in the NE was increased after F-actin depletion by LatB (Fig. 7D, E). This result provides additional evidence that MLKS2 binds to F-actin in an ARM domain-dependent manner.

**Figure 7.**
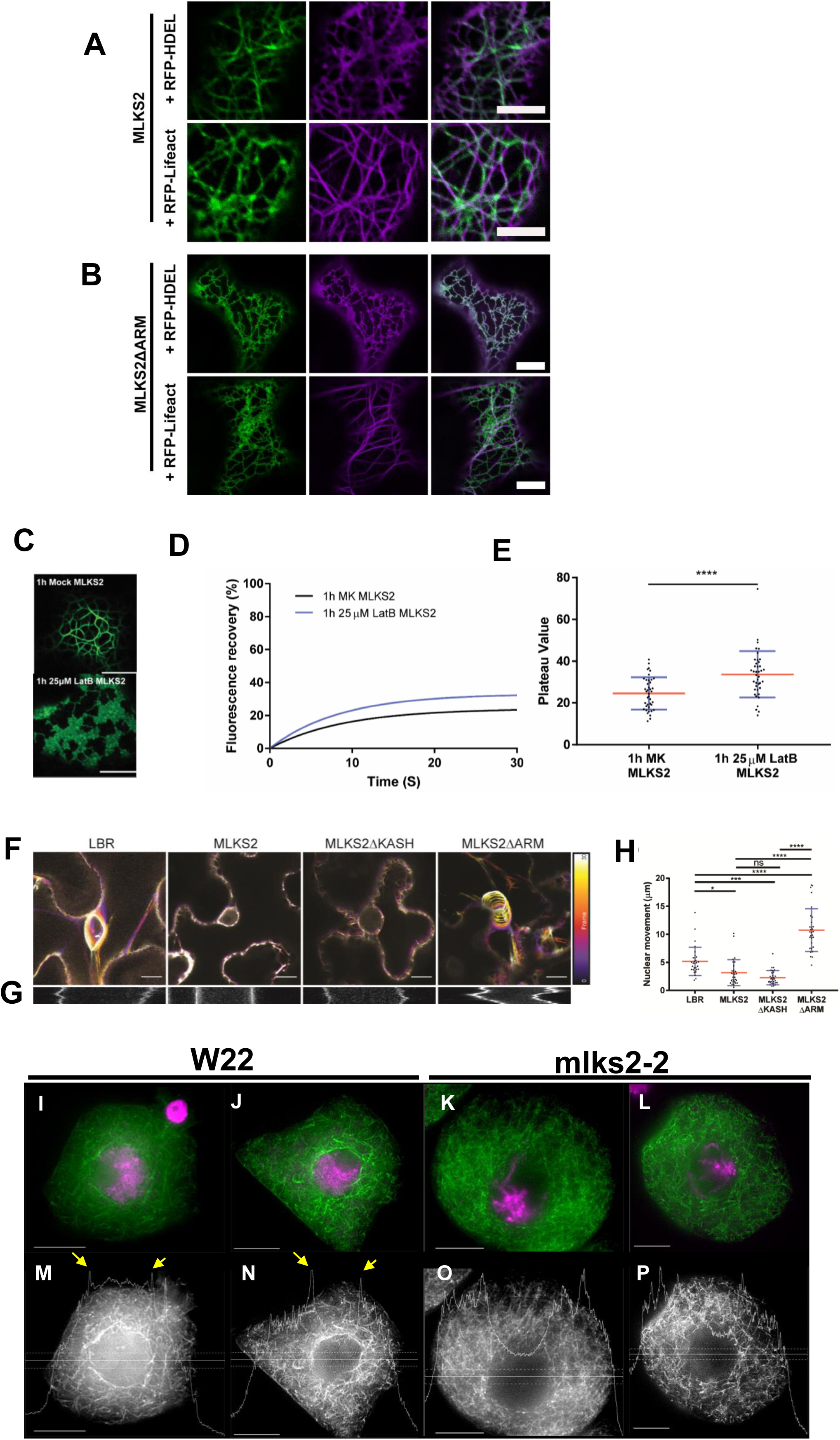
MLKS2 interaction with actin. Co-localization of ER marker (RFP-HDEL, magenta) or actin marker (RFP-LifeAct, magenta) with A) GFP-MLKS2 or B) GFP-MLKS2ΔARM. C) GFP-MLKS2 localization in F-actin-depleted cells (25 uM LatB) phenocopies GFP-MLKS2ΔARM with an ER-like staining pattern. D) Depolymerization of the actin cytoskeleton with LatB results in increased GFP-MLKS2 FRAP recovery compared to mock (MK) controls. E) FRAP Plateau value of MLKS2 mock (MK) and LatB treated as shown in D). For whisker plots, blue lines denote SD error bars, red lines denote mean. Student’s T-test used to test statistical significance. **** = P≤0.0001. Scale bar denotes 10 µm. F) Temporal color-coded projections of randomly selected LBR, MLKS2, MLKS2ΔKASH and MLKS2ΔARM nuclei imaged every 10 seconds over 5 minutes. G) Kymographs of nuclear movement shown in F) for a different nucleus. H) Quantification of total nuclear movement over time, imaged for LBR, MLKS2, MLKS2ΔKASH and MLKS2ΔARM. N = at least 30 nuclei imaged across three experimental repeats per treatment; ANOVA statistical test used. Ns = P≥0.05, * = P≤0.05, *** = P≤0.001 and **** = P≤0.0001. (I-L) DAPI (magenta) and phalloidin (green) stained early prophase maize meiocytes. (M-P) Gray-scale images of phalloidin staining of cells shown in panels I-L. Line trace plots show intensity of phalloidin in the middle of the cell marked by horizontal band, illustrating the spike in perinuclear actin in wild-type but not mutant nuclei. Scale bar denotes 15 µm.

Given these results together with the previously established roles for some KASH proteins’ actin-based nuclear anchoring (Gundersen and Worman, 2013; Huelsmann and Brown, 2014; Zhou et al., 2014), we next asked if GFP-MLKS2 could affect nuclear mobility in a live cell assay. The nuclear mobility baseline was first established in *N. benthamiana* using GFP-LBR (Graumann et al., 2010) as a general NE marker (Fig. 7F-H). For these experiments, nuclear movement was tracked in live cells for five min by imaging at 10 s intervals to obtain total distance migrated. Compared to the GFP-LBR-expressing control, the nuclei expressing GFP-MLKS2 were dramatically immobilized as shown by temporal color-coded projections, kymograph analysis (Fig. 7F-G), and measurements of total distance of nuclear movement over the 30-frame time-lapse (Fig. 7H). Expression of MLKS2 and MLKS2ΔKASH resulted in significant decreases in nuclear movement compared to the LBR control. In contrast, the nuclear anchoring activity of MLKS2 could be abolished by deletion of the ARM domains (Fig. 7H). These live nuclear mobility assays define an ARM-dependent nuclear anchoring activity for MLKS2, consistent with the core function of the LINC complex in connecting the nucleus to the cytoskeleton.

Given the role of the NE dynamics in meiotic prophase and our findings that MLKS2 interacts with F-actin in somatic cells, we returned to maize to check genetic evidence of MLKS2-actin interactions in pollen mother cells at meiotic prophase (Fig. 7I-P). The overall appearance of phalloidin-stained F-actin in W22 was similar to that of *mlks2-2* meiocytes, with two notable exceptions. First, the F-actin in the mutant plants appeared to exhibit a subtle phenotype of somewhat discontinuous or fragmented F-actin structures (compare fiber lengths of Fig. 7 panels J and L versus N and P). Secondly, and most importantly, we observed that the *mlks2-2* meiocytes exhibited a conspicuous loss of perinuclear F-actin staining. Perinuclear actin is common in maize meiocytes (Murphy et al., 2014) and intensity line profile tracings show clear evidence of F-actin concentration spikes around the nucleus (yellow arrows, Fig. 7M, N). However, we were unable to observe by inspection (Fig. 7I-L) or line profile analysis (Fig. 7M-P) any such perinuclear actin staining in *mlks2-2* mutant meiocytes. These findings implicate MLKS2 in the formation of perinuclear F-actin to achieve proper meiotic chromosome behavior.

### MLKS2 can rescue Arabidopsis triple *wip* nuclear shape phenotype

Finally, the functional conservation of MLKS2 in plants was addressed using a genetic rescue assay. Previously, it has been shown that deletion of certain KASH proteins, such as AtWIP proteins, causes nuclei to become more circular (Zhou et al., 2012). GFP-MLKS2 was expressed in Arabidopsis harboring mutations in three *WIP* genes to determine if the maize SINE-group KASH protein could correct the nuclear shape defect phenotype. For this experiment, stable transformants expressing GFP-MLKS2 were grown and nuclei were imaged and analyzed from two different tissues, as shown in Figure 8. Circularity index measurements of leaf and root nuclei showed that maize GFP-MLKS2 complemented the nuclear shape phenotype of the triple-*WIP* mutant in Arabidopsis by restoring the nuclei to wild-type length in two different cell types. These experiments demonstrate that a GFP-tagged maize SINE-family KASH protein can substitute for Arabidopsis *WIP*-family KASH proteins in nuclear shape control. In summary, the findings from this study establish that the maize KASH protein MLKS2 plays a role in multiple biological pathways including stomatal complex development, meiotic nuclear architecture and positioning, meiotic chromosome segregation, and production of viable pollen.

**Figure 8.**
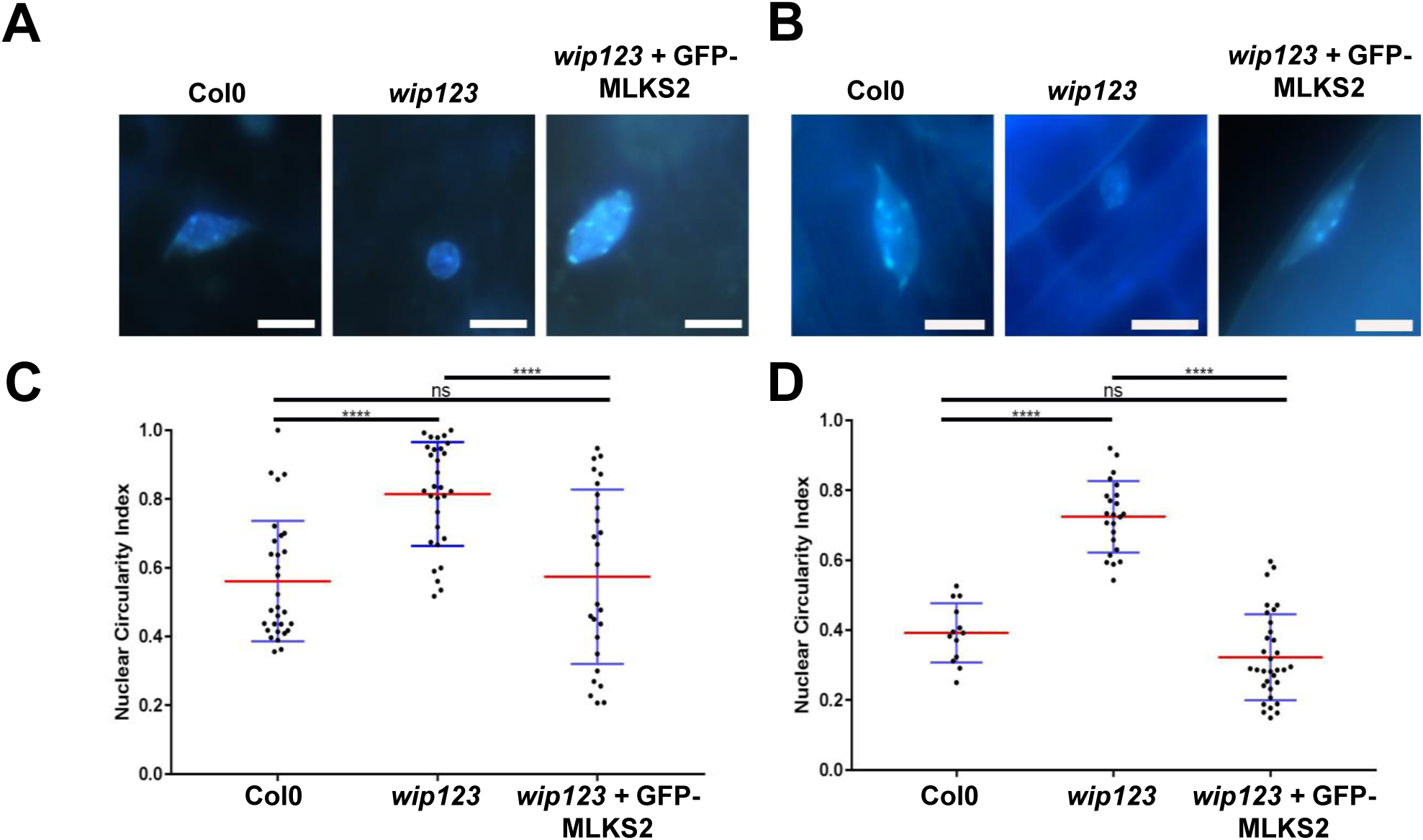
Arabidopsis triple *wip* mutant phenotype rescued with MLKS2. DAPI stained representative images of A) leaf and B) root nuclei from three Arabidopsis genotypes: *Arabidopsis thaliana* WT (Col0); (Columbia WT strain), *wip123* (*AtWIP*-type KASH triple mutant where *wip123* refers to genotype *wip1-1*, *wip2-1*, *wip3-*(Zhou et al., 2015b), or *wip123* plus GFP-MLKS2 (WIP triple mutant transformed with GFP-MLKS2). (C, D) Nuclear circularity index summaries for these same tissues and genotypes are shown below each tissue/genotype combination, where 1 is a perfectly round nucleus and values lower than 1 are a measure of nuclear elongation. Arabidopsis WIP triple mutant nuclei are significantly more rounded (leaf CI=0.81±0.03; root CI=0.72±0.02) than WT (leaf CI=0.56±0.03; root CI=0.39±0.02) whereas GFP-MLKS2 complemented nuclei are similar to WT (leaf CI=0.52±0.03, root CI=0.32±0.02); p>0.05=ns, p<0.0001=****.

## Discussion

In this study we characterized a core LINC component, MLKS2, in maize using genetic and cellular assays. We showed that MLKS2 localizes to two cellular compartments, the NE as expected and cytoplasmic ER where it colocalizes with actin via the ARM domain region. Genetic analysis uncovered mutant phenotypes in both vegetative and reproductive organs while documenting the first case of a plant KASH required for proper telomere clustering and chromosome behavior in meiotic prophase. Interactions of MLKS2 with F-actin via the ARM domains of MLKS2 may represent a general mechanistic principle that explains how MLKS2 might control nuclear shape and positioning. The pleiotropic nature of the mutants and MLKS2’s deep conservation in plants may be important for a variety of tissue-specific biological functions that require specialized control of nuclear morphology and dynamics. For instance, MLKS2 may play a role in female meiosis and ovule development where highly organized nuclear positioning occurs in the mature ovary, pollen development and nuclear migration of sperm and vegetative nuclei, and fertilization itself. Interestingly, the overall plant morphology of both the wild type (WT) and homozygous mutant plants was largely normal.

At the onset of this study, the diagnostic ARM domains of the SINE group of plant KASH and previous findings from Arabidopsis AtSINE1 (Zhou et al., 2014) pointed to a possible MLKS2 interaction with F-actin. Interestingly, GFP-AtSINE1, but not GFP-AtSINE2, colocalizes with F-actin when expressed in tobacco or Arabidopsis (Zhou et al., 2014). Medicago SINE-group KASH genes, *MtSINE1* and *MtSINE2*, also encode proteins that colocalize with F-actin (Newman-Griffis et al., 2019). The ARM domains alone may not, therefore, be predictive of F-actin binding. Consistent with this is the fact that a variety of proteins have ARM domains, but their binding partners vary (Coates, 2003). In this study, there exists a clear preponderance of evidence, direct and indirect, that MLKS2 has F-actin binding activities. In the heterologous expression system, MLKS2 localizes to the NE, as expected for a KASH domain protein, and to a cytoskeletal-like reticulate network that colocalizes with F-actin and ER markers at the cell periphery (Fig. 7A). Two complementary cell localization experiments, latrunculin-B depletion of F-actin and ARM domain deletions, uncover a possible cytoplasmic function that does not involve the nucleus, but rather deploys MLKS2 to tether the ER to cytoplasmic F-actin. In this regard, MLKS2 resembles members of the Networked (NET) proteins, as defined by Hussey and colleagues (Deeks et al., 2012). Although MLKS2 does appear to fit the definition of membrane-bound plant-specific actin-binding adapter protein, it lacks the tryptophan-rich NET actin-binding (NAB) domain sequence, common to NET families 1-4 (Hawkins et al., 2014). The NET NAB domain is thought to confer actin-binding activity, whereas for MLKS2 that activity may come from the ARM domain. We note that the non-nuclear cytoplasmic MLKS2 may result from overexpression or compromised targeting in the heterologous expression system used. It nonetheless predicts that misregulation by overexpression may present cellular dysfunction phenotypes generally associated with the cytoplasm and specifically associated with the ER and cortical F-actin.

The actin interaction with core LINC NE components provides mechanistic insights into several cellular functions of MLKS2 in both vegetative and reproductive parts of the maize plant. We focused on three major nuclear phenotypes assayed primarily in somatic cells: (1) positioning within the cell, (2) shape, and (3) mobility. Nuclear positioning was implicated by genetic analysis of stomatal complex development, nuclear shape was examined via rescue of root hair nuclei rounding up, and nuclear mobility was directly measured in live cells.

Nuclear positioning defines cell polarity. One well-characterized example of highly polarized cells associated with asymmetric cell divisions is the normal stomatal complex development in leaves. In wild-type maize, guard mother cells (GMC) signal to adjacent subsidiary mother cells (SMC) to commence a series of cellular events involving nuclear migrations and localized actin patches that coordinate the eventual production of a mature stomatal complex (Müller et al., 2009). The mature stomatal complex is characterized by two guard cells flanked by subsidiary cells, displaying a bilateral symmetry. Mutations known to disrupt stomatal complex development in maize include the *discordia* (*dcd1*), the *panglass* (*pan1*, *pan2*), and the *brick* (*brk1*, *brk2*, and *brk3*) mutants (Facette et al., 2015). The brick proteins are part of the SCAR complex which interacts with ARP to form actin patches on the GMC side of the SMC. How the nucleus moves towards the actin patches is not known. The KASH phenotypes reported here bear a striking resemblance to those of *pan1* and *brk1*, placing *MLKS2* in the stomatal complex development pathway. Given that MLKS2 ARM domains appear to interact with F-actin, it is tempting to speculate that the developmental defect seen in *mlks2* reflects a failure to coordinate nuclear positioning with the cortical-ER-patches known to occur at the region adjacent to the inducing GMC (Giannoutsou et al., 2011).

Nuclear shape maintenance is one of the fundamental functions performed by LINC complex proteins. Nuclear morphology is commonly altered in LINC mutants, some of which present pathological conditions such as envelopathies (Burke and Stewart, 2006; Janin et al., 2017). In eudicot plants, disruption of LINC components cause alterations in root hair nuclear shape phenotypes (Newman-Griffis et al., 2019; Oda and Fukuda, 2011; Tamura et al., 2013; Zhou et al., 2015a). In maize, we similarly observed that *mlks2* nuclei change to more compact and spherical shapes than those from normal cells (Fig. 5A-C). There is growing evidence that disruption of any component of the core LINC components or even the LINC-interacting nucleoskeletal proteins can cause a “rounded up” nuclear phenotype. The finding that maize GFP-MLKS2 was able to rescue the nuclear shape phenotype in Arabidopsis plants lacking three WIP proteins is important for two reasons. First, it represents the first demonstration of cross-species genetic rescue of a LINC component. Second, it suggests that even diverged LINC components may possess some degree of structural redundancy in establishing a balance of forces required for maintaining nuclear shape. The triple *WIP* rescue by a maize SINE-group KASH may reflect, therefore, the capacity of MLKS2 to replace a NE-cytoskeletal tether normally formed by the WIP--WIT-- MyosinXI-i-actin bridge (Tamura et al., 2013).

Nuclear mobility in plants is associated with a wide variety of cellular processes including symbiosis, pathogenesis, fertilization, and division planes in development (Griffis et al., 2014). Nuclear mobility can either be inferred from variable positioning in fixed cells, such as the eccentric nucleus stage of early meiotic prophase (Fig. 4), or by direct time-lapse tracking in live cells (Fig. 7F-H). Here we observed both. Our results show that the mobility of tobacco nuclei is reduced or stopped by heterologous expression of GFP-MLKS2, and this immobilization is reversed by removing the ARM domain region of MLKS2. These findings, consistent with our genetic analyses, implicate MLKS2 in nuclear anchoring, which may further explain, for example, the appearance of wild-type elongated nuclei in root hair cells (Fig. 5). The lack of perinuclear actin around the meiotic nucleus in *mlks2-2* plants further suggests that MLKS2 may function to bind or recruit filamentous actin to the perinuclear cytoplasm. In this way, MLKS2 may play an organizational or mechanical role whereby microfilaments, microtubules, or their associated motor proteins may direct the dynamics of meiotic nuclei (Sheehan and Pawlowski, 2009).

The role of the NE in meiotic chromosome behavior has been recognized for over a century. In recent decades, the LINC complex has been shown to be required for telomere clustering at the bouquet stage, as first described in fission yeast (Chikashige et al., 2009). Here we show that MLKS2 is required for normal bouquet formation and proper chromosome segregation. Our findings represent the first genetic evidence that a plant KASH interacts with the actin cytoskeleton during male meiosis. These results suggest that actin may be more important in plant meiosis than previously recognized. This idea is consistent with live imaging of maize meiotic prophase chromosome and nuclear movements, both of which were stopped following MF depletion by latrunculin-B treatment (Sheehan and Pawlowski, 2009). In our mutants, the earliest cytological meiotic defect is seen at the zygotene stage, when the telomere bouquet is normally formed. The *mlks2* mutants exhibit defects in telomere clustering and nuclear positioning. These early stage problems may later manifest in the failure of homologs to synapse and recombine with high fidelity. MLKS2 may also have post-bouquet functions such as interlock resolution and telomere dispersal, ideas consistent with detection of irregular and pronounced clumping at late prophase (Fig. 4G-J). The chromosome segregation laggards, micronuclei, and pollen sterility phenotypes may therefore reflect multiple upstream failures of disomic pairing and recombination. Interestingly, several of our *mlks2* phenotypes resemble those described for mutations in AtSUN genes (Varas et al., 2015) or the meiotic kinesin1-like gene *AtPSS1 (Duroc et al., 2014)*. *AtSUN1-1* and *AtSUN2* double mutants also show partial polarization of telomeres, absence of full synapsis, presence of unresolved interlock-like structures, and reduction in the mean cell chiasma frequency which manifests in the formation of laggard chromosomes during anaphase and severely reduced fertility (Varas et al 2015). Several of these phenotypes are the same or similar to what we observed with *mlks2*, which is not surprising given that SUN and KASH are core components of the LINC complex. Similarly, mutations in *AtPSS1* leads to phenotypes shared with mlks2, including production of univalents and aneuploid cells after meiosis (Duroc et al 2014). Findings such as these, together with the large body of evidence implicating microtubules in plant meiotic chromosome behavior (Cowan and Cande, 2002; Cowan et al., 2002; Driscoll and Darvey, 1970; Higgins et al., 2016; Shamina, 2003; Thomas and Kaltsikes, 1977) suggest that meiotic chromosome behavior likely requires both actin and tubulin cytoskeletal systems. Indeed, the LINC complex in general, and MLKS2 in particular, may provide a coordinating structure that integrates cytoskeletal systems to affect nuclear architecture.

In conclusion, this study identifies MLKS2 as a core LINC component with multiple functions in fundamental plant vegetative and reproductive processes. Analysis of the ARM domain sheds light on how the SINE group of plant KASH proteins can interact with the actin cytoskeleton. In Figure 9, we present a working model for MLKS2 where we propose that MLKS2 interacts with F-actin via its N-terminal ARM domains in wild-type (Fig. 9A) but not mutant plants (Fig. 9B). The MLKS2 protein could extend a Velcro-type hook into the cytoplasm to bind generally or specifically with cytoskeletal linear polymers, most likely F-actin. This interaction between MLKS2 and F-actin could be achieved by direct or indirect contact (Fig. 9B, compare left and right). In tobacco (Fig. 9C,D), the expression of GFP-MLKS2 or GFP-MLKS2ΔKASH results in nuclear anchorage according to kymograph analysis, suggesting the ARM domain can anchor the nucleus even if the KASH domain is absent. The anchorage is relieved by removal of the ARM domains (Fig. 9D, right half). The models based on heterologous expression in *N. benthamiana* include GFP-MLKS2 in the core LINC complex via a presumed interaction with tobacco SUN (NbSUN, grey), or a demonstrated interaction with maize SUN2 (red, ZmSUN2). These findings highlight the importance of investigating individual components of the LINC complex in order to understand how the NE coordinates cytoplasmic and nuclear phenomena to carry out fundamental biological processes at the cellular level.

**Figure 9.**
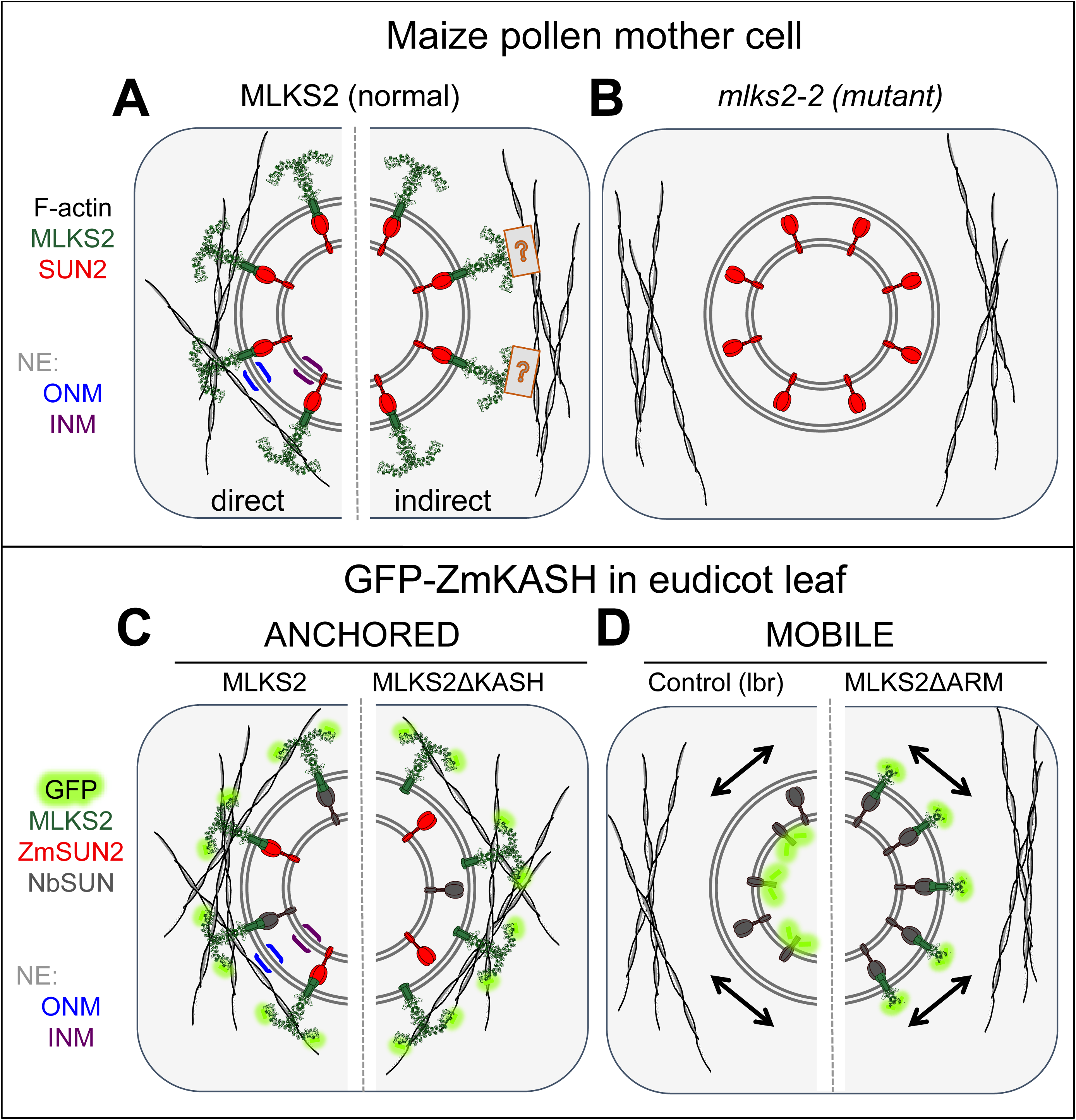
Summary diagrams and models of ZmMLKS2. Summary diagram illustrates how MLKS2 may interact with F-actin to produce the genetic and heterologous expression phenotypes reported in this study. A) MLKS2 is presumed to be arranged with an ARM domain-containing alpha solenoid structure in the cytoplasm, where it interacts with F-actin directly (left half) or indirectly through a hypothetical connector depicted by the boxed question mark (right half). B) The *mks2-2* mutant has lost the ability to bind or contribute to the recruitment of F-actin. Not depicted here are data from MLKS2 in vegetative (leaf, root) organs. Results from kymograph analysis of GFP-MLKS2 expressed in *N. benthamiana* are summarized for experiments that showed C) anchored nuclei in cells expressing full length MLKS2 or MLKS2ΔKASH, or D) mobile nuclei in cells expressing a control NE marker (lbr) or MLKSΔARM. In these diagrams, evidence for interaction with ZmSUN2 co-expression is indicated (red) on the basis of FRAP assays or depicted from presumed interactions with tobacco SUN, NbSUN (grey). In these diagrams (C,D), the MLKS2 interaction with F-actin is depicted as direct for convenience, but the alternative indirect mode (A, right half) of MLKS2 interaction with F-actin remains a formal possibility.

## Materials and Methods

### Microscopy with tobacco and Arabidopsis

*Nicotiana benthamiana* (*Nb*) transformation was performed as described (Sparkes et al., 2006). Agrobacterium cultures were used at an OD of 0.05 and plants imaged 3 days post-infiltration. All imaging was performed on a Zeiss LSM 800 confocal system equipped with a 100X 1.4NA lens. Bi-directional scanning was used for all imaging. For DAPI excitation the 405nm laser was used and emission collected at 400-491 nm. For GFP excitation the 488 nm laser was used and emission collected between 491-617 nm. For mCherry excitation the 561 nm laser was used and emission collected from 561-620 nm. Colocalization imaging was performed with multi-track line switching to avoid fluorescence bleedthrough, and 4x frame averaging was used. For FRAP acquisition a 5X digital zoom was used with a frame time of 0.15s. Fluorescence recovery was monitored by consecutive imaging of 240 frames. Bleaching was performed after 5 control frames were imaged using the 488 nm laser at 100X power for 20 iterations in a 160×160 pixel ROI containing the NE. For nuclear movement assays, nuclei were imaged at 10 s intervals for 30 frames (5 minutes). We used LBR-GFP as a control for baseline nuclear movement as described in (Graumann et al., 2010). For DAPI staining, *N. benthamiana* or *A. thaliana* tissue was incubated in 5µg/ml DAPI and 0.001% Triton X-100 for 10 min before imaging.

For image analysis *ZEN* (Blue edition, v2.3.69.1000, Zeiss) was used and figures compiled in *Illustrator* (v22.1, Adobe). For FRAP analysis, normalized intensity values were calculated as described (Martinière et al., 2012), and one-phase association for each replicate and combined curves plotted in *GraphPad* v7.04. For nuclear movement assays, manual tracking was used to track the center of nuclear movement for each experimental condition, and FIJI (Image J version 1.52h) was used to generate temporal color-coded projections and kymographs over a 25 µm line. All whisker plots showing data points were generated in *GraphPad* v7.04. Student’s T-test or one-way ANOVA statistical tests were used depending on the suitability of the data, as indicated in each figure legend. The number of replicates used for each experiment can be found in the figure legends. Circularity index analysis was performed as described (Zhou et al., 2012).

### Gene constructs

Gene constructs, nomenclature, and sequence information for the clones used in this study are listed in Table S1. GFP-MLKS2 constructs were synthesized (GenScript Biotech Corp.) using the full-length ORF of MLKS2 fused to the C-terminal end of eGFP-FLAG-HA. The synthetic clone (GFP-MLKS2) was obtained in a pUC18 vector, inserted at the *Bam*HI restriction site to produce a plasmid designated pHWBF07. From this vector, the eGFP-FLAG-HA-MLKS2 sequence was PCR amplified with *att* flanking primers (Table S2) using high fidelity Q5 polymerase (NEB) and cloned into pDONR221 vector by BP cloning (Cat. No. 1235019, Invitrogen) to generate the entry clone designated pHWBF07EC. The eGFP-FLAG-HA-MLKS2 sequence from the entry clone was transferred to vector pH7WG2 (Karimi et al., 2002) by Gateway LR recombination (Cat. No. 1235019, Invitrogen) to obtain the expression vector designated pPK1Fexp. Deletion constructs were generated from the full-length entry clone as a template to PCR-amplify required regions with internal primers containing flanking *att* sequences and cloned into pDONR211 by Gateway BP cloning method. These deletion constructs were also subcloned into pH7WG2 destination vector by Gateway LR recombination. Full length and domain deletion constructs for ZmSUN2 were generated as described in (Gumber et al., 2019).

### Maize plant material and genotyping

The transposon insertion alleles of *MLKS2* (unique insertion IDs: mu1038603 and mu1058535) were identified through MaizeGDB (http://www.maizegdb.org/) and derived from the UniformMu transposon mutagenesis project (McCarty and Meeley, 2009; Settles et al., 2007). Seeds for stock number UF-Mu04133 carrying allele *mlks2-1* or stock number UF-Mu07312 carrying allele *mlks2-2* were obtained from the Maize Genetics Cooperation Stock Center (http://maizecoop.cropsci.uiuc.edu/). For “wild-type” maize referred to here as W22, we used color-converted W22 obtained from Hugo Dooner (Waksman Inst., Rutgers, New Jersey, USA) derived by Brink (Brink, 1956). The seeds were planted at the Florida State University Mission Road Research Facility (Tallahassee, FL, USA) and propagated by out-crossing to W22 in summer 2014. In the fall, the progeny seeds were grown in a greenhouse in the King Life Sciences building (Biological Science Dept, Florida State University, Tallahassee, FL). The segregating outcross plants were self-crossed to obtain mutant plants from among the progeny.

For PCR genotyping, leaf samples from 4-week-old maize W22 and plants from families segregating for MLKS2 alleles were harvested, frozen, and stored at −80 C. Genomic DNA was isolated using a modified mini-CTAB method from approximately 2 cm^2^ leaf tissue in 96 well plates in a Mixer Mill (Retsch®) as described (Labonne et al., 2013). Plants were genotyped using three different primer pair combinations to accurately predict the genotype as described by (McCarty et al., 2013). For the -*mlks2-1* allele, DNA extracted from each of the UF-Mu04133 stock plants was individually amplified with 603F1-603R1 gene specific primer pair, as well as a gene specific primer in combination with the Tir6 primer corresponding to terminal inverted repeats of the Mu1 transposable element (603F1-Tir6 and 603R1-Tir6). Similarly, for the *mlks2-2* allele, DNA extracted from each of the UF-Mu07312 stock plants was individually amplified with 535F2-535R2, 535F2-Tir6 and 535R2-Tir6 primer pairs. Normal W22 DNA was also amplified with all of these primer pairs. All the primer sequences used for genotyping are listed in Table S3. The PCR amplification products were resolved by agarose gel electrophoresis followed by ethidium bromide staining. The PCR products were sequence-verified using TA cloning in pCR™4Blunt-TOPO® Vector (Invitrogen cat # K2875-20). M13F and M13R vector primers were used for sequencing (Molecular Cloning Facility, Department of Biological Sciences, Florida State University) and the resulting sequences were aligned with the W22 reference genome sequence to validate the mu transposon element insertion sites.

### RNA extraction and qRT-PCR

After one round of outcrossing to W22 and two rounds of self crossing, seeds were grown in the greenhouse to harvest meiotic tassels from V13 -stage plants. The tassels were frozen in liquid nitrogen and stored at −80°C until further use. The tissue was ground in liquid nitrogen and 100 mg of the resulting powder used for RNA extraction as per manufacturer’s instructions (Qiagen RNeasy kit, Cat # 74904). The extracted RNA was treated with amplification-grade DNase I (ThermoFisher, Cat # 18068015) for 15 min at room temperature, then treated with RNA clean and concentrator-25 kit (Zymo Research, Cat # R1017). The resulting RNA was inspected for quantity and quality by UV spectroscopy and gel electrophoresis, respectively. Superscript III reverse transcriptase (ThermoFisher, Cat# 18080093) was used to reverse transcribe 1µg of high quality RNA into cDNA according to the manufacturer’s instructions. The resulting cDNA was used as a template to amplify ∼ 100 bp products downstream of Mu1 insertion sites in *mlks2-1* and *mlks2-2* using three gene-specific primers. Two housekeeping genes CDK1 and RPN2 were amplified as internal controls (Lin et al., 2014). The products were amplified using SYBR Green Master Mix (Applied Biosystems) on a 7500 Fast Real-Time PCR System (Applied Biosystems). Relative expression change was calculated using the 2ΔΔCt method (Livak and Schmittgen, 2001). Two technical replicates were carried out to ascertain the accuracy of the procedure.

### Microscopy in maize

For 3D telomere FISH analysis, meiotic stage tassels were harvested from W22 and *mlks2-2* plants grown in the greenhouse. Flowers were microdissected and fixed in 1X Buffer A with 4% paraformaldehyde for 1 hour on rotary shaker at room temperature as described (Bass et al., 1997). The fixed anthers were microdissected and embedded in acrylamide pads as described in (Howe et al., 2013) and subjected to telomere FISH using a 0.13 µM fluorescein-conjugated telomere-specific oligonucleotide probe, MTLF. The meiocytes were counterstained with 3 µg/mL DAPI then mounted in vectashield (Vector labs) and imaged using a 3D-deconvolution microscope equipped with a 60X lens and CCD camera (Deltavision, GE Healthcare). Telomere coordinates in X, Y, and Z were identified using the 3D MODEL tool of SoftWorx program (DeltaVision, GE Healthcare) and pairwise telomere distances were calculated using R software for Euclidean distances.

For actin staining, meiotic-staged anthers were fixed in PHEMS buffer supplemented with 8% paraformaldehyde for 2 hours with shaking at room temperature as described (Chan and Zacheus Cande, 1998). The fixed anthers were microdissected on a glass slide in PBS (pH 7.0) and stained with 3.3 µM Alexa Fluor™ 594 Phalloidin (Thermo Fisher Scientific) and DAPI (1.5 µg/mL) for 20 min at room temperature in the dark. The cells were imaged using a Deltavision 3D deconvolution microscope.

For pollen viability, anthers from male flowers just before dehiscence were fixed in Carnoy’s fixative (6 alcohol:3 chloroform:1 acetic acid) for a minimum of 2 hours at room temperature. The fixed anthers were placed on glass slides and stained with modified Alexander’s stain containing Malachite green (0.01%), Acid Fuchsin (0.05%) and Orange G (0.005%) as described (Peterson et al., 2010) to differentiate viable (magenta) pollen grains from aborted (green) pollen grains. Bright field images of pollen grains were collected on Revolve microscope (Echo Labs). At least 300 pollen grains each from 3 plants of every genotype were counted to calculate pollen viability.

For maize root hair imaging, seeds were sterilized with 1% bleach, 0.01% Triton X-100 in deionized water for 10 min, rinsed and soaked in water overnight. The next day seeds were spread on wet paper towels, kept dark and moist for 5-6 days, then roots were cut off and fixed in Buffer A (Howe et al., 2013) plus 4% final concentration paraformaldehyde for 1 hour, with rotation at room temperature. Small pieces of roots with root hair from the elongation zone were placed on glass slides and chopped to small pieces with a razor blade. The tissue was stained with 3 µg/mL DAPI for 20 min at room temperature, mounted with VECTASHIELD, and imaged on an EVOS fluorescence microscope (Thermo Fisher Scientific). The images were processed using the *Analyze particle* function of *ImageJ* to measure the longest diameter and circularity of the nuclei.

For leaf and stomatal complex imaging, plants were grown in the greenhouse and the 4th leaf was harvested at its first appearance. For developing stomatal complex, up to 2 cm tissue from the base of the leaf was harvested, and for mature stomatal complex, the remainder of the leaf was harvested and fixed in FAA fixative (3.7%, 10% Formalin, 50% ethanol and 5% acetic acid in water) for at least 15 min at room temperature as described in (Facette et al., 2015). The leaf tissue was cut in thin longitudinal strips and treated with PBS supplemented with 1% Triton for 10 min to permeabilize the tissue. The tissue was then rinsed and incubated in a solution of propidium iodide (Life technologies) and DAPI for 30 min. The tissue was rinsed with PBS to get rid for excess dye and mounted in water. The tissue was imaged on a 3D deconvolution microscope at 60X magnification.

## List of abbreviations

FRAP: Fluorescence recovery after photobleaching
DPI: Days post infiltration
OD: Optical density
MLKS2: Maize LINC KASH AtSINE-like2
LINC: Linker of nucleoskeleton and cytoskeleton
NE: Nuclear envelope
INM: Inner nuclear membrane
ONM: Outer nuclear membrane

---

**Figure S1.**
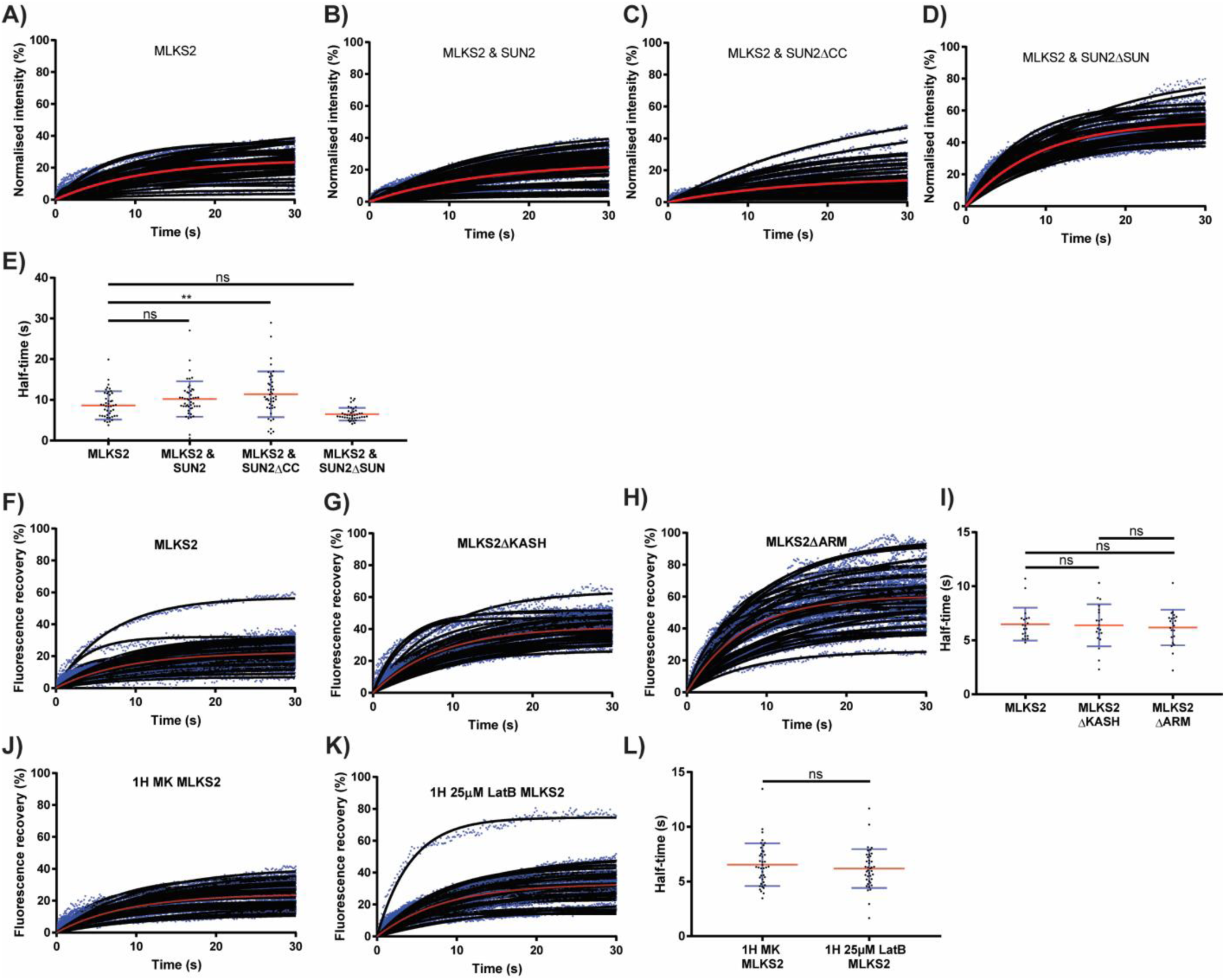
Individual nucleus data from FRAP experiments presented throughout the main figures. A) FRAP recovery curves from all individual nuclei imaged with GFP-MLKS2 for comparison to co-expression with SUN2. B) FRAP recovery curves from all individual nuclei imaged with GFP-MLKS2 and co-expressing mCherry-SUN2. C) FRAP recovery curves from all individual nuclei imaged with GFP-MLKS2 and co-expressing mCherry-SUN2ΔCC. D) FRAP recovery curves from all individual nuclei imaged with GFP-MLKS2 and co-expressing mCherry-SUN2ΔSUN. E) Halftime values from FRAP recovery experiments shown in A-D. F) FRAP recovery curves from all individual nuclei imaged with GFP-MLKS2 as control during GFP-MLKS2ΔKASH and GFP-MLKS2ΔARM experiments. G) FRAP recovery curves from all individual nuclei imaged with GFP-MLKS2ΔKASH. H) FRAP recovery curves from all individual nuclei imaged with GFP-MLKS2ΔARM. I) Halftime values from FRAP recovery experiments shown in F-H. J) FRAP recovery curves from all individual nuclei imaged with GFP-MLKS2 after 1 hour mock treatment, for comparison to LatB treatment. K) FRAP recovery curves from all individual nuclei imaged with GFP-MLKS2 after 1 hour 25µM LatB treatment. L) Halftime values from FRAP recovery experiments shown in J and K. Recovery curves blue dots show individual data points, black lines individual fluorescence recovery curves, and red curves average recovery curves for all nuclei. For whisker plots, blue lines denote error bars showing standard deviation, red lines show mean values. Statistical tests performed were either one-way ANOVA (E and I) or Student’s t-test (L) as appropriate. ns = P≥0.05, ** = P = ≤ 0.01.

**Figure S2.**
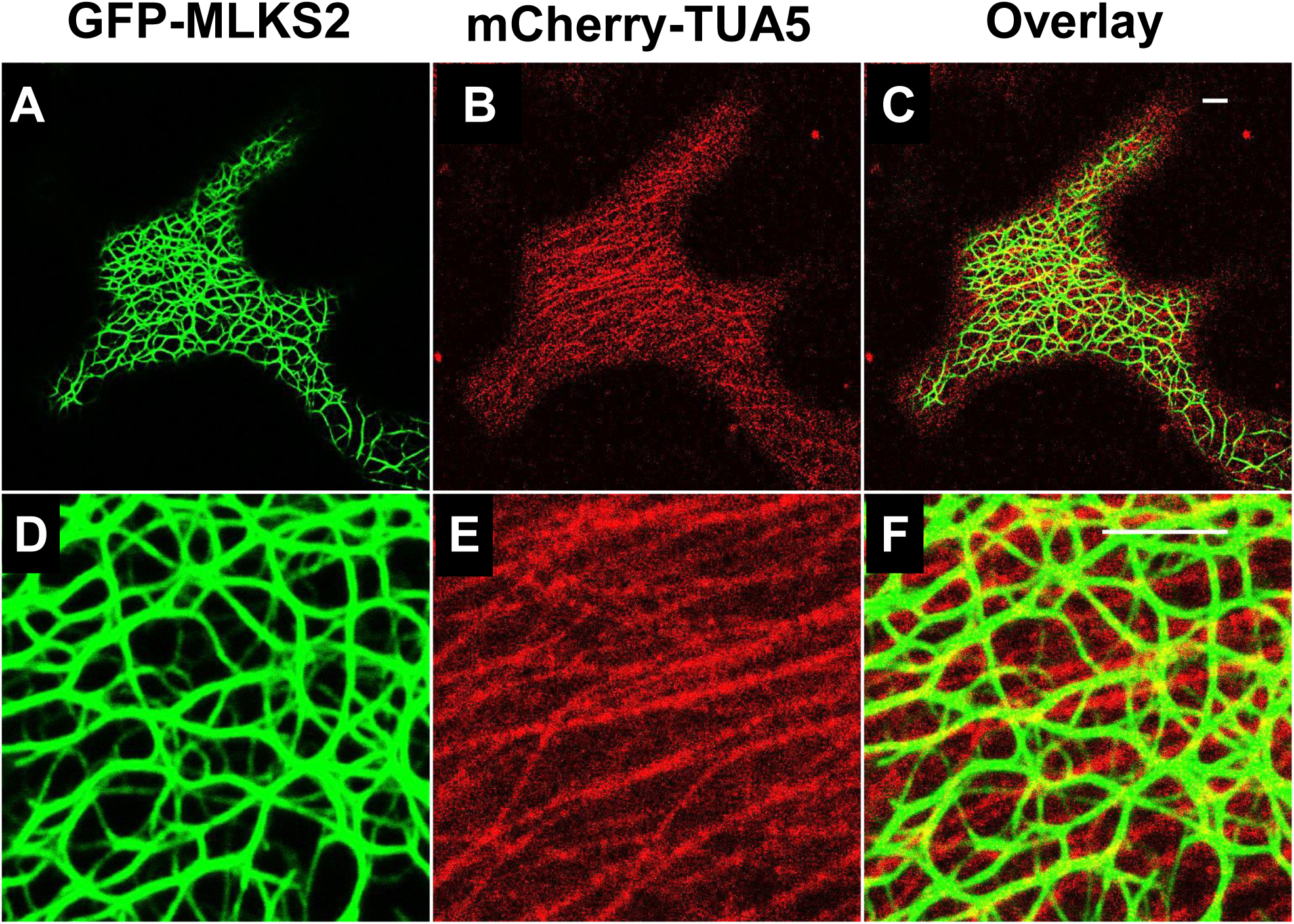
Co-expression of GFP-MLKS2 and tubulin alpha-5 (TUA5) Representative images of *N. benthamiana* leaves transiently co-transformed with GFP-MLKS2 and mcherry-TUA5. MLKS2 and microtubules do not seem to colocalize. Scale bar denotes 10 µm.

**Table S1.**
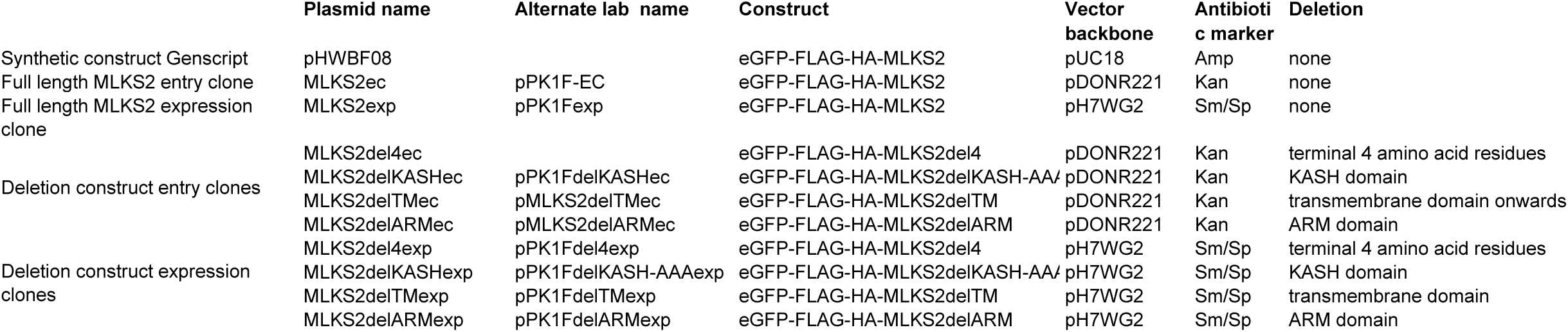
Plasmid information (Gumber et al.)

|  | Plasmid name | Alternate lab name | Construct | Vector backbone | Antibiotic marker | Deletion |
| --- | --- | --- | --- | --- | --- | --- |
| Synthetic construct Genscript | pHWBF08 |  | eGFP-FLAG-HA-MLKS2 | pUC18 | Amp | none |
| Full length MLKS2 entry clone | MLKS2ec | pPK1F-EC | eGFP-FLAG-HA-MLKS2 | pDONR221 | Kan | none |
| Full length MLKS2 expression clone | MLKS2exp | pPK1Fexp | eGFP-FLAG-HA-MLKS2 | pH7WG2 | Sm/Sp | none |
| Deletion construct entry clones | MLKS2del4ec |  | eGFP-FLAG-HA-MLKS2del4 | pDONR221 | Kan | terminal 4 amino acid residues |
|  | MLKS2delKASHec | pPK1FdelKASHec | eGFP-FLAG-HA-MLKS2delKASH-AA/ | pDONR221 | Kan | KASH domain |
|  | MLKS2delTMec | pMLKS2delTMec | eGFP-FLAG-HA-MLKS2delTM | pDONR221 | Kan | transmembrane domain onwards |
|  | MLKS2delARMec | pMLKS2delARMec | eGFP-FLAG-HA-MLKS2delARM | pDONR221 | Kan | ARM domain |
| Deletion construct expression clones | MLKS2del4exp | pPK1Fdel4exp | eGFP-FLAG-HA-MLKS2del4 | pH7WG2 | Sm/Sp | terminal 4 amino acid residues |
|  | MLKS2delKASHexp | pPK1FdelKASH-AAAexp | eGFP-FLAG-HA-MLKS2delKASH-AA/ | pH7WG2 | Sm/Sp | KASH domain |
|  | MLKS2delTMexp | pPK1FdelTMexp | eGFP-FLAG-HA-MLKS2delTM | pH7WG2 | Sm/Sp | transmembrane domain |
|  | MLKS2delARMexp | pPK1FdelARMexp | eGFP-FLAG-HA-MLKS2delARM | pH7WG2 | Sm/Sp | ARM domain |

**Table S2.**
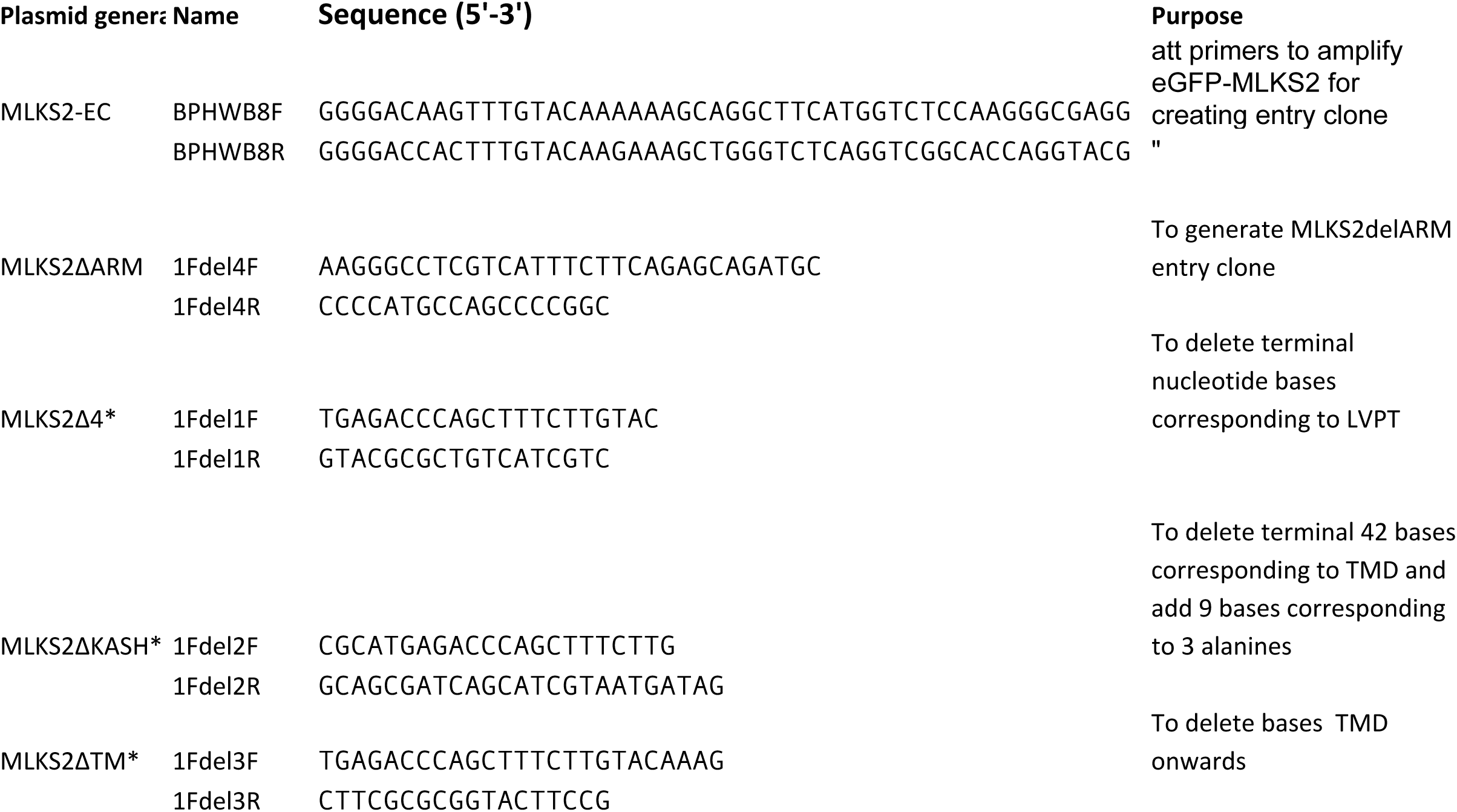
List of primers used for construct cloning (Gumber et al.)

**Table S3.**
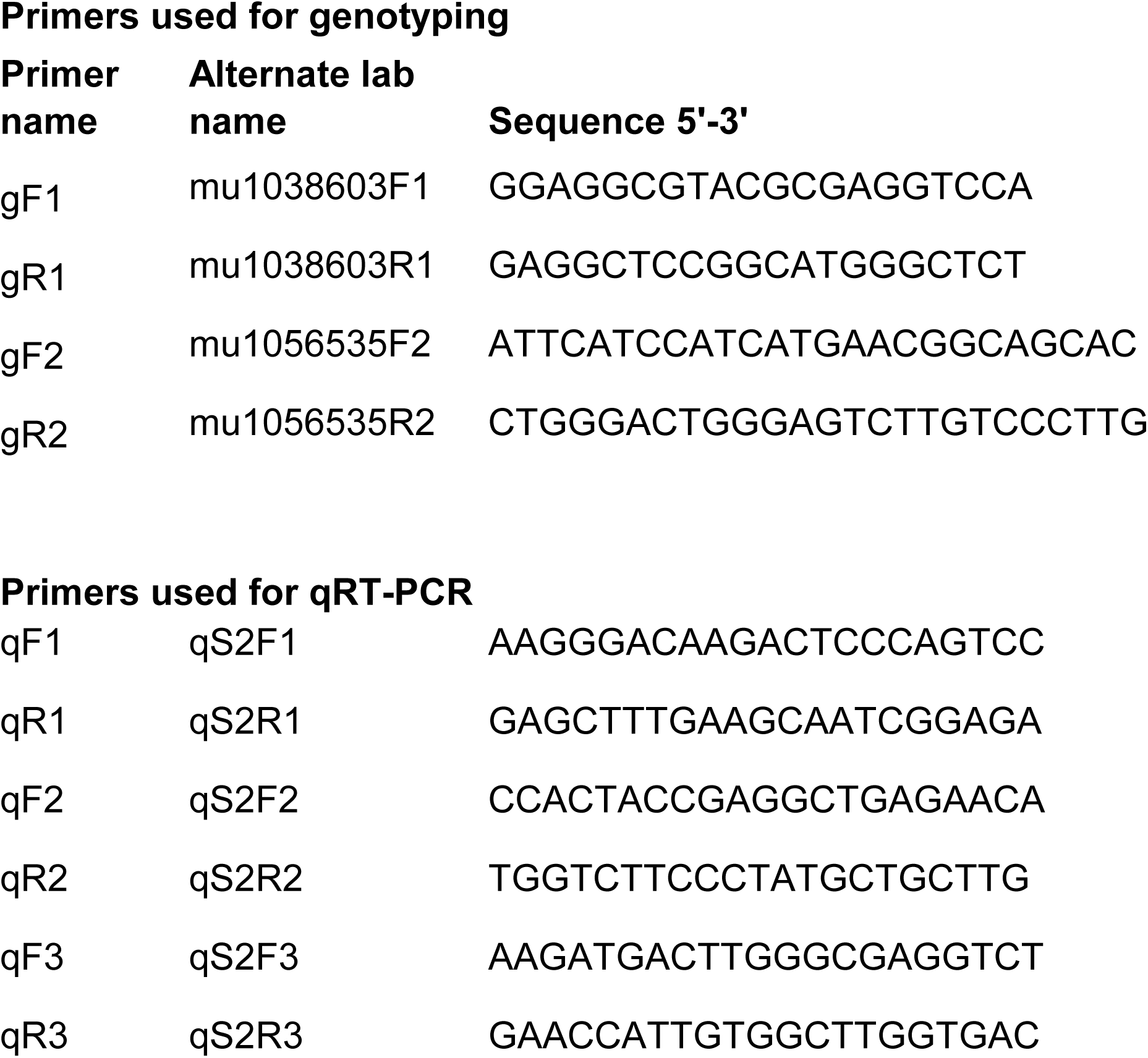
List of genotyping primers (Gumber et al.)

